# Oxytocin facilitates social behavior of female rats via selective modulation of interneurons in the medial prefrontal cortex

**DOI:** 10.1101/2024.07.15.603543

**Authors:** Stephanie Schimmer, Alan Kania, Arthur Lefevre, Konstantinos Afordakos, Kai-Yi Wang, Julia Lebedeva, Andrey Rozov, Androniki Raftogianni, Rishika Tiwari, Shai Netser, Ana Zovko, Huma Shaheen, Jonas Schimmer, Ryan Patwell, Clémence Denis, Valentin Grelot, Hugues Petitjean, Lan Geng, Dimitri Hefter, Arjen Boender, Yuval Podpecan, Franziska Schommer, Tim Schubert, Anna Sanetra, Aleksandra Trenk, Anna Gugula, Rene Hurlemann, Shlomo Wagner, Yulong Li, Ferdinand Althammer, Anna Blasiak, Sarah Melzer, Hannah Monyer, Alexandre Charlet, Marina Eliava, Valery Grinevich

## Abstract

The hypothalamic neuropeptide oxytocin is best known for its prosocial behavioral effects. However, the precise anatomical and cellular targets for oxytocin in the cortex during social behavior remain elusive. Here we show that oxytocin neurons project directly to the medial prefrontal cortex where evoked axonal oxytocin release facilitates social behaviors in adult female rats. In conjunction, we report that local oxytocin receptor-expressing (OTR^+^) cells are predominantly interneurons, whose activation promotes social interaction. Notably, this prosocial effect persists even under physiological challenge (hunger), pointing to a dedicated prosocial circuit capable of overriding primary survival drives. We further demonstrate that activation of these OTR^+^ interneurons inhibits principal cells specifically projecting to the basolateral amygdala, thus providing a putative mechanism of selective oxytocin action in this sociability-promoting cortical network.

## INTRODUCTION

The neuropeptide oxytocin (OT) plays a central role in modulating various types of social behavior and is primarily synthesized in the hypothalamic paraventricular (PVN), supraoptic (SON), and accessory nuclei (Swanson and Sawchenko 1983, Grinevich, Knobloch-Bollmann et al. 2016). Hypothalamic OT is subsequently transported to the posterior lobe of the pituitary gland and released into the blood where it elicits milk ejection during lactation and uterine contractions during childbirth (Russell, Leng and Douglas 2003). Beyond its peripheral effects, OT is actively involved in modulating neural circuits within the brain. Particularly, direct social interaction strongly activates the OT system via OT neurons in the PVN (Tang, Benusiglio et al. 2020). Axonal collaterals of OT neurons extend to various forebrain regions (Knobloch, Charlet et al. 2012), which coincides with the expression of OT receptors (OTRs) at the site of putative neuropeptide release (Chini, Verhage and Grinevich 2017). These areas, including the central nucleus of the amygdala, nucleus accumbens, ventromedial hypothalamic nucleus, lateral septum, and hippocampus (Marlin and Froemke 2016, Grinevich and Stoop 2018), are pivotal in processing social information, emotional responses, and stress regulation (Veenema and Neumann 2008). The infusion of OT, evoked OT release, or activation of OTR neurons in these regions trigger a broad spectrum of region-specific social behaviors, such as social fear (Knobloch, Charlet et al. 2012, Neumann and Slattery 2015, Menon, Grund et al. 2018), social transmission and contingency (Ferretti, Maltese et al. 2019), social memory (Gur, Tendler and Wagner 2014), social avoidance (Osakada, Yan et al. 2024), and aggression (Calcagnoli, Stubbendorff et al. 2014). While most studies show that OT’s behavioral effects arise from subcortical circuits, emerging work has started to uncover its role in cortical signaling. OT signaling in the murine auditory cortex and its role in nursing, has been extensively studied by Froemke and colleagues (Marlin, Mitre et al. 2015, Valtcheva and Froemke 2018, Schiavo, Valtcheva et al. 2020). Moreover, OT has been shown to drive top-down modulation of early sensory processing in the olfactory bulb to enhance social recognition (Oettl, Ravi et al. 2016). Additionally, several studies have directly implicated cortical OT signaling in the regulation of socio-sexual behaviors: in the medial prefrontal cortex (mPFC), Heintz and colleagues identified OTR-expressing (OTR^+^) interneurons that gate female sociosexual behavior and revealed a sex-dimorphic OTR–CRH interaction shaping social and anxiety-related responses (Nakajima, Görlich and Heintz 2014, Li, Nakajima et al. 2016). Moreover, OTR-expressing glutamatergic neurons in the mPFC have been shown to selectively regulate social recognition memory (Tan, Singhal et al. 2019).

The mPFC is instrumental in orchestrating various higher order and computational aspects of social behavior in mammals, including but not limited to social motivation (Lee, Rhim et al. 2016), recognition of social cues (Gifford III, MacLean et al. 2005, Levy, Tamir et al. 2019), and decision-making (Kingsbury, Huang et al. 2019). The connections of the mPFC to other brain areas like the amygdala (Myers-Schulz and Koenigs 2012) and hypothalamus (Reppucci and Petrovich 2016) suggest its role in top-down executive control, such as guiding goal-directed social behaviors. In rodents, the mPFC can be divided into three distinct brain subregions, each with unique functions and connectivity: the infralimbic cortex (ILC), the prelimbic cortex (PrL) and the cingulate cortex (Cgt) (Heidbreder and Groenewegen 2003). Considering the widespread influence of OT across various brain regions and its known input to the mPFC (Knobloch, Charlet et al. 2012), it is likely that OT modulates mPFC functions.

Despite the inspiring demonstrations that OT action and/or OTR^+^ neurons in the mPFC modulate paternal (He, Young et al. 2019) and sexual behavior (Nakajima, Görlich and Heintz 2014) in voles and mice, respectively, the OT-modulated circuit that mediates same-sex affiliative (non-sexual) social behavior in the mPFC remains largely unexplored. To address this gap, we aimed to describe the pattern of OT innervation in the mPFC (1), the composition of the local OT receptive cells (2), and their downstream target structures (3), which might promote social behavior in adult female rats.

## RESULTS

### Evoked OT release in the infralimbic cortex facilitates social behavior in female rats

Immunohistochemical (IHC) labelling revealed the presence of OT projections through the entire mPFC **(Figure 1 A, B1-2)**. The highest density of OT axons was found in the infralimbic cortex (ILC) compared to the cingulate (Cgt) and prelimbic (PrL) cortices (49.9±9.2 vs. 1.5±0.8 axonal segments per mm^2^ in 50µm coronal slices; this and following data are presented as mean±sd; p=0.0002, details of statistical tests are provided in **Supplemental Document 1**; sample sizes are reported in the figure legends) **(Figure 1 B3)**. Based on this finding, we focused specifically on the ILC in all follow-up experiments. Considering that the IHC approach labelled all OT fibers independent of origin, we next employed a viral approach to label specific nuclei in the hypothalamus to determine the source of OT inputs to the ILC. Therefore, we injected a recombinant adeno-associated virus (rAAV) equipped with the OT promoter (rAAV-OTp-Venus) either in the PVN or the SON of female rats (**Figure 1 C1)**, and analyzed fiber density in the ILC with a semi-automated Imaris approach. We found significantly denser innervation originating from the SON than from the PVN (135.4±65.6 vs. 51.8±32.1 10^3^ x µm^3^ sum volume of fibers, p=0.0253) (**Figure 1 C2**). For both hypothalamic nuclei we found that they mostly innervate layer 5/6, with additional substantial innervation in layer 1, while the least innervation was found in layer 2/3 (**Figure 1 C3**) (p=0.0007 for layer 1 vs. layer 5/6 SON projections, p<0.0001 for layer 2/3 vs. layer 5/6 SON projections; significance of post hoc tests of ANOVAs and mixed effect models are indicated with # in figures).

**Figure 1.**
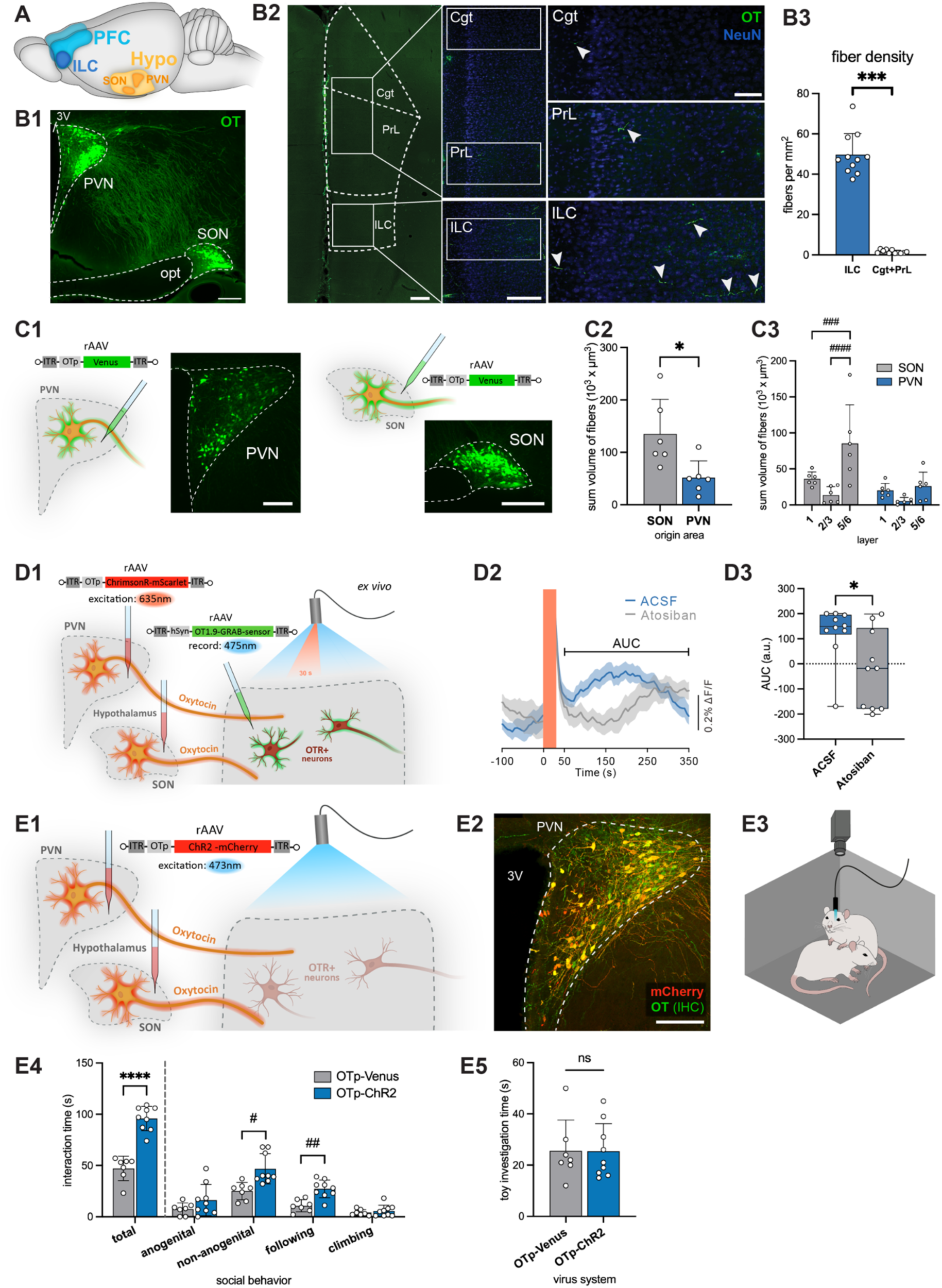
Evoked OT release in the ILC facilitates social behavior in female rats. **A** The rat brain regions of interest: PVN and SON in the hypothalamus and the ILC as part of the mPFC. **B** OT innervation in the medial prefrontal cortex. **B1** Immunohistochemical staining for OT in the hypothalamus showing OTergic nuclei, the PVN and SON. 3V: third ventricle, opt: optic tract. Scalebar 500µm. **B2** Coronal section of the mPFC subdivisions (Cgt, PrL, ILC) and high-magnification images of OT fiber distribution for each sub region. Scalebar left: 500µm, middle: 300µm, right 100µm. **B3** Density of OT-positive fibers quantified in digitized images. The number of OT-positive fibers per mm^2^ was significantly higher in the ILC, compared to the mPFC dorsal subdivisions (n=11 sections obtained from 5 rats). **C** Innervation from the PVN and SON. **C1** Injection scheme depicting the injection of a rAAV expressing Venus under the control of the OT promoter (rAAV-OTp-Venus) into either the PVN or SON in separate animals, with respective injection sites. Scalebar 250µm. **C2** Automated analysis of fibers in the ILC quantifying their density by the sum of their volume originating from the PVN or SON reveals stronger innervation from the SON. **C3** Localized analysis of fiber innervation originating from the PVN or SON depending on the layer in the ILC (layer 1, layer 2/3 or layer 5/6) reveals most fibers in layer 5/6 (n=6 sections obtained from 3 animals per group). **D** Axonal OT release in the ILC *ex vivo*. **D1** Injection scheme depicting the injection of a rAAV expressing the excitatory ChrimsonR under the control of the OT promoter (rAAV-OTp-ChrimsonR-mScarlet) into the hypothalamic nuclei of female rats and injection of a rAAV expressing an OT-sensor (rAAV-OT1.9-GRAB-sensor) in the ILC. Stimulation of the OT fibers in the ILC (625 nm light, 30 Hz, 10 ms light on, for 30s). **D2** Peri-stimulus GRAB-OT1.9 fluorescence changes in the ILC in ACSF-only solution (blue) or in the presence of Atosiban (OTR-antagonist, gray). Orange square marks the stimulation period. Bracket indicates the timeframe for area under the curve (AUC) analysis in panel D3. Data presented as mean±sem. **D3** Recordings in ACSF condition showed higher AUC, calculated between 50 and 350 seconds after light stimulation, compared to recordings in the Atosiban condition (n=10 sections). **E** Optogenetic activation of OT axons in the ILC during free social interaction. **E1** Injection scheme depicting the injection of a rAAV expressing the excitatory ChR2 under the control of the OT promoter (rAAV-OTp-ChR2-mCherry) into the hypothalamic nuclei of female rats and implantation of an optic fiber above the ILC. **E2** Injection site showing high specificity of the rAAV-OTp-ChR2-mCherry (red) after counterstaining for OT (green). Scalebar 200µm. **E3** 5-minute social interaction paradigm with an unknown conspecific; blue light (BL) stimulation present for the first two minutes of the session. **E4** Animals injected with OTp-ChR2 (n=9 animals) show significantly increased social interaction time compared to control animals (injected with OTp-Venus, n=7 animals), especially increased non-anogenital sniffing and following behavior. **E5** Investigation time of a toy rat in the open field remains unaltered by optogenetic activation. Statistical significance is indicated as * p<0.05, ** p<0.01, *** p<0.001, **** p<0.0001. Statistical significance of post-hoc tests is indicated as # p<0.05, ## p<0.01, ### p<0.001, #### p<0.0001. Error bars show standard deviation. For details on statistical tests please refer to Supplemental Document 1.

Next, we investigated whether OT axons release the neuropeptide in the ILC. To this end, we injected a rAAV expressing an opsin under the control of the OT promoter into the PVN and SON (rAAV-OTp-ChrimsonR-mScarlet, **Extended Data Figure S1A, right:** patch clamp verification of opsin functionality, **Extended Data Figure S1 B1:** representative injection site). In parallel, we injected a rAAV expressing the GRAB OT 1.9 sensor (rAAV-hSyn-OT1.9-GRAB) (kindly provided by Prof. Yulong Li, GRAB OT 1.0 published in: Qian, Wang et al. 2023) in the ILC, and performed imaging experiments in acute ILC slice preparations. We observed a significant increase of GRAB OT 1.9 fluorescence emission upon stimulation of OT fibers (625 nm light, 30 Hz, 10 ms light on, for 30s) **(Figure 1 D1),** consistent with OT release and consecutive binding (Iwasaki, Lefevre et al. 2023). This effect was blocked by incubation of the tissue with the OT receptor antagonist (Atosiban) **(Figure 1 D2-3, Extended Data Figure S1 B2-3)** (123.4±110.5 vs. -23.9±155.8 AUC, p=0.0232).

To functionally probe the hypothalamic-ILC OT pathway, we next expressed channelrhodopsin 2 (ChR2) in OT neurons of the PVN and SON (rAAV-OTp-ChR2-mCherry, **Extended Data Figure S1A left** for patch clamp verification of opsin functionality) and implanted an optic fiber above the ILC **(Figure 1 E1-2)**. Next, rats were exposed to an unknown, age-, strain-and sex-matched conspecific **(Figure 1 E3)** in an open field arena (OF) for 5 minutes, where we analyzed the social behavior throughout the session upon light stimulation of OT fibers in the ILC for the first 2 minutes of the session. In keeping with previous studies (Knobloch, Charlet et al. 2012, Tang, Benusiglio et al. 2020), we employed adult female rats to exclude the aggressive component of social behavior often observed between unfamiliar males (Schuster, Berger and Swanson 1988). Social interaction time was significantly increased (total interaction time 47.1±11.8s vs. 95.9±11.9s, p<0.0001) during and immediately after optogenetic stimulation with blue light (BL, 470 nm, 30 Hz,10 ms pulses, laser power ∼10 mW, 120 s) compared to controls expressing Venus fluorescent protein instead of ChR2. This effect was mainly due to significantly increased non-anogenital sniffing and following behaviors (25.0±8.5s vs. 46.8±14.7s, p=0.0103 and 10.7±5.9s vs. 27.2±8.6s, p=0.0019, respectively) **(Figure 1 E4)**.

Importantly, no difference was observed between the two groups when comparing the time they spent investigating a toy rat (25.6±12.0s vs. 25.4±10.8s, p=0.9826) **(Figure 1 E5)**, indicating that the behavior reflects a social component rather than general interest and novelty seeking. No changes in locomotor activity, exploratory, or anxiety-related behaviors were found **(Extended Data Figure S1 C)**. Altogether, these results indicate that optogenetic activation of the OTergic hypothalamic-ILC projections are sufficient to enhance social behavior in female rats.

### OTR^+^ neurons in the ILC are intrinsically active during social behavior

Next, to target and manipulate OTR^+^ neurons in the ILC we utilized newly generated OTR-Cre knock-in rats (Iwasaki, Lefevre et al. 2023). The rats were injected with a rAAV expressing GFP in a Cre-dependent (double-floxed-inverted-orientation: DIO) manner into the ILC (rAAV-Ef1a-DIO-GFP) **(Figure 2 A1,** see also **Extended Data Figure S2 B** for further characterization of the rat line and protein expression). Semi-automated quantification revealed that among the entire ILC neuronal population, visualized by NeuN **(Figure 2 A2)**, 1.2% of neurons expressed OTRs with a higher occurrence in superficial layers 2/3 compared to the deeper layers 5/6 (1.6% and 0.9%, respectively, p<0.0001) **(Figure 2 A3, Extended Data Figure S2 A)**. To monitor OTR^+^ neuronal intrinsic activity, we performed *in vivo* fiber photometry during social interaction utilizing a rAAV expressing a Cre-dependent Ca^2+^ sensor (rAAV-Ef1a-DIO-GCaMP6s) and implanted optic fibers **(Figure 2 B1-2)**. During repeated 5-minute social interaction sessions, multiple behavioral events (non-anogenital and anogenital sniffing, and investigation of a toy) were scored and the aligned calcium activity of OTR^+^ neurons was compared **(Figure 2 B3)**. Social sniffing behaviors performed by the test rat were accompanied by a significant increase of calcium signal in OTR^+^ neurons compared to baseline (z score for anogenital: 0.12±0.01, non-anogenital: 0.18±0.01), while toy investigation did not lead to a significant change in calcium signal (0.04±0.01) (mean±sem, p<0.0001 for anogenital and non-anogenital sniffing compared to baseline, p=0.4707 for toy) **(Figure 2 B4)**. Furthermore, we also compared the amplitude of calcium signals during social events that were initiated by the test rat (active social behavior) with the amplitude of calcium signals during social events performed by the stimulus rat (received social behavior) **(Figure 2 B5**). Only active social behavior increased calcium-dependent fluorescence in OTR^+^ neurons in the ILC (z score for active: 0.12±0.006, p<0.0001, received: -0,01±0.008, p=0.9035, mean±sem) compared to baseline **(Figure 2 B6)**. Therefore, we conclude that OTR^+^ neurons in the ILC are highly and specifically active during active social behavior.

**Figure 2.**
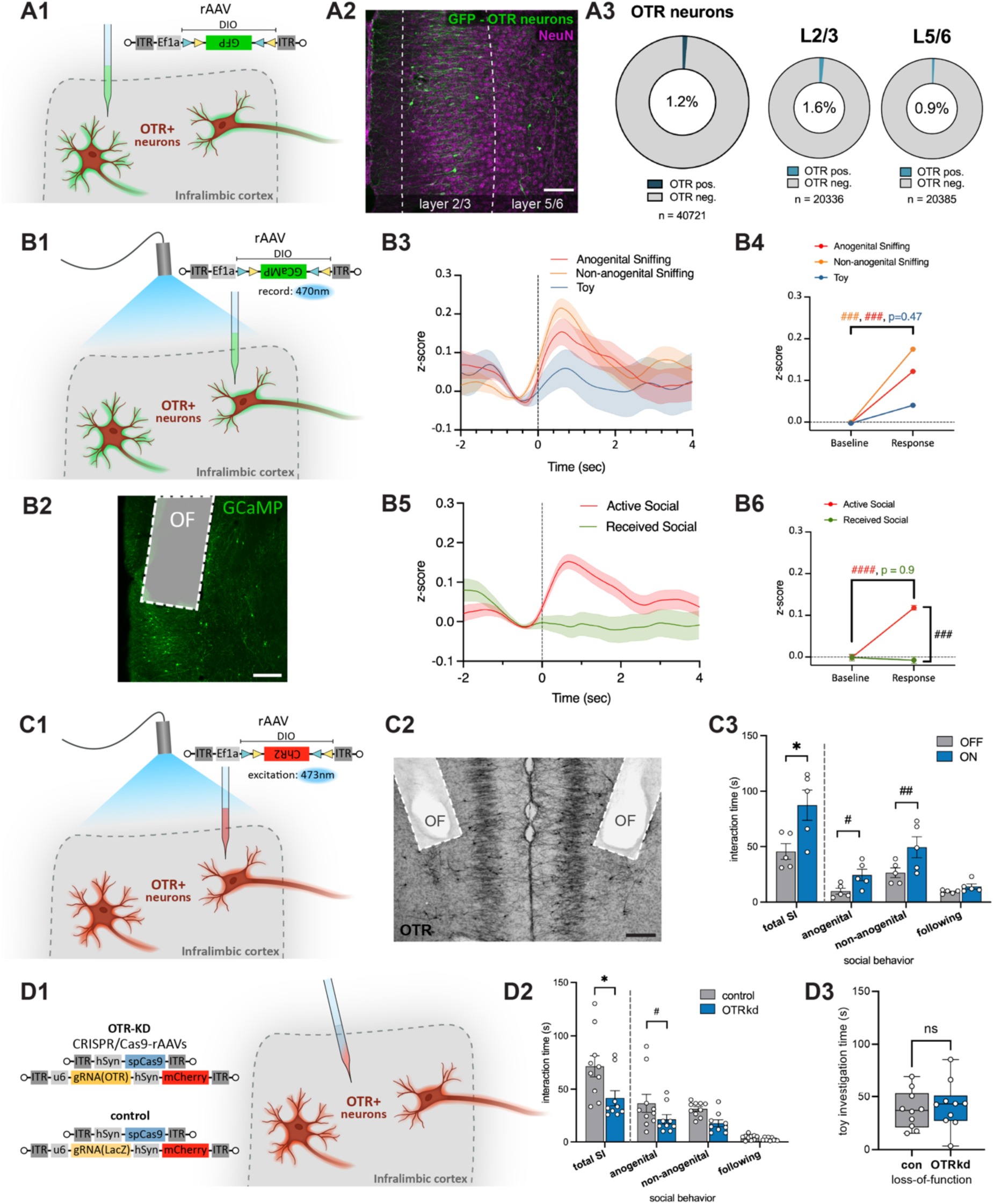
OTR neurons in the ILC are active during social behavior and modulate sociability. **A** OTR^+^ neurons in the ILC. **A1** Injection scheme depicting the injection of a rAAV expressing GFP in a Cre-dependent manner and **A2** counterstaining for NeuN. Scalebar 150µm. **A3** Semi-automated, high-throughput analysis of ILC slices reveals that 1.2% of neurons in the ILC express OTRs, with very similar distributions in the superficial layers 2/3 and deeper layers 5/6 (n=11 sections obtained from 4 animals; n in the figure indicate the number of tracked neurons). **B** Recording of OTR^+^ neurons during social interaction. **B1** Injection scheme depicting the injection of a rAAV expressing GCaMP6 in a Cre-dependent manner and implantation of an optic fiber into the ILC of OTR-Cre-rats to record Ca^2+^ activity from OTR^+^ neurons. **B2** Injection site showing OTR^+^ neurons infected with GCaMP (green) and the optic fiber (OF) tract. Scalebar 200µm. **B3** 1321 observations of neuronal activity during different social and non-social behaviors collected during several social interaction sessions in multiple animals (n=4) were aligned to the start of the behavior (t=0) and averaged per behavior (anogenital sniffing (n=391 events), non-anogenital sniffing (n=706 events), toy investigation (n=224 events)). **B4** The strongest increase in intracellular calcium in OTR^+^ neurons in the ILC was observed during anogenital and non-anogenital sniffing behavior, while the calcium activity was not significantly increased during toy investigation compared to baseline activity. **B5** Neuronal calcium activity during either active (n=1368 events) or received (n=600 events) social interactions. **B6** OTR^+^ neuron calcium activity in the ILC was increased only during active social behaviors, but not during received social behaviors compared to baseline activity. **C** Optogenetic activation of OTR^+^ neurons in the ILC during social interaction. **C1** Injection scheme depicting the injection of rAAV expressing the excitatory optogenetic channel Channelrhodopsin (ChR2) in a Cre-dependent manner into the ILC of female rats and implantation of an optic fiber above the ILC. **C2** Injection site showing OTR^+^ neurons infected with ChR2 (black) and the optic fiber tract. Scalebar 200µm. **C3** Animals spend significantly more time socially interacting if the BL is ON in the first two minutes of the session, compared to when it is OFF (n=5 animals). This effect is significantly driven by anogenital and non-anogenital sniffing. **D** CRISPR/Cas9-mediated OTR depletion decreases social interaction time. **D1** Injection scheme depicting the injection of a virus system which utilizes CRISPR/Cas9 technology to specifically and functionally knockdown OTRs locally in the ILC (n=9 animals loss-of-function group, n=10 animals control group). **D2** OTR loss-of-function significantly decreases social interaction (SI) time compared to control animals, especially for non-anogenital sniffing behavior. **D3** Investigation time of a toy rat in the open field remains unaltered by OTR loss-of-function. Statistical significance is indicated as * p<0.05, ** p<0.01, *** p<0.001, **** p<0.0001. Statistical significance of post-hoc tests is indicated as # p<0.05, ## p<0.01, ### p<0.001, #### p<0.0001. Error bars show standard deviation. For details on statistical tests please refer to Supplemental Document 1.

### OTR^+^ neurons in the ILC facilitate social behavior

After demonstrating that OTR^+^ neurons in the ILC are active during social interaction, we aimed to artificially activate them using optogenetic stimulation, in order to test whether activation of ILC OTR^+^ neurons affects social behavior. For that, we injected a rAAV expressing ChR2 in a Cre-dependent manner (rAAV-Ef1a-DIO-ChR2) into the ILC of OTR-Cre-rats and implanted optic fibers above the injection sites **(Figure 2 C1-2)**. During the first 2 minutes of a 5-minute social interaction session, OTR^+^ neurons were either activated by BL (30 Hz,10 ms pulses), or no light was applied for a baseline social behavior readout. Compared to the baseline, BL activation significantly increased social interaction times (total interaction time 45.6±15.8s vs. 87.4±30.6s, p= 0.0124), and more specifically increased anogenital and non-anogenital sniffing (9.8±5.9s vs. 24.4±11.6s, p= 0.0426 and 26.4±9.7s vs. 49.4±21.2s, p=0.0036, respectively) **(Figure 2 C3)**.

To verify that the endogenous OT/OTR^+^ signaling in the ILC modulates sociability, we introduced a local and discreet mutation in the OTR gene (Boender, Boon et al. 2023) **(Extended Data Figure S2D)** by injecting a mixture of two rAAVs expressing Cas9 (rAAV-hSyn-spCas9) and a guide RNA specific for the OTR-sequence (rAAV-u6-gRNA(OTR)-hSyn-mCherry) or LacZ-sequence which served as a control (rAAV-u6-gRNA(lacZ)-hSyn-mCherry) **(Figure 2 D1)**. Injection sites and viral expression were verified post mortem **(Extended Data Figure S2C)**. We first confirmed the depletion of OTRs in the ILC functionally by bath application of the selective OTR agonist, TGOT, and *ex vivo* patch clamp electrophysiological recordings of OTR^+^ neurons in both groups **(Extended Data Figure S2E)**. TGOT-induced membrane depolarization was significantly abolished by the viral OTR depletion compared to the control group **(Extended Data Figure S2F)**. Notably, the ILC OTR depletion was characterized by a significantly decreased social interaction time (total social interaction time 71.5±31.1s vs. 41.3±22.2s, p=0.0259) with an unfamiliar conspecific. This effect was driven by a significant decrease in anogenital sniffing (36.2±26.6 vs. 21.2±13.0s, p=0.0493), while non-anogenital sniffing also trended towards a decrease in time **(Figure 2 D2)**. Importantly, no difference was observed regarding the investigation time of a toy rat (37.7±18.0s vs. 41.7±22.5s, p=0.6656) **(Figure 2 D3)**.

### OTRs are predominantly expressed in GABAergic neurons of the ILC

A multiplex fluorescent in situ hybridization (RNAscope) approach was used to characterize the types of ILC OTR^+^ neurons **(Figure 3 A1-3)**. Probes for both OTR as well as Cre were used to verify the specificity of our OTR-Cre rat line; whereas probes for vGlut1 (vesicular glutamate transporter), vGlut2 and vGAT1 (vesicular GABA transporter) to identify both glutamatergic and GABAergic neurons. We observed that 92.6±8.6% of Cre-positive neurons were also OTR-positive, and vice versa 82.9±8.5% of OTR-positive neurons were also Cre-positive, confirming high specificity of the transgenic OTR-Cre line **(Figure 3 A4)**. Notably, the vast majority of OTR-positive neurons co-expressed vGAT1 (87.63%), while only a small fraction co-expressed vGlut1 (9.14%), and none co-expressed vGlut2 **(Figure 3 A5)**. Moreover, we observed differences between the superficial and deeper layers of the ILC: OTR^+^ neurons in layer 2/3 were predominantly GABAergic (96.84% vGAT1-positive), while OTR^+^ neurons in layer 5/6 were prevalently GABAergic (78.02% vGAT1-positive), but also included a substantial number of glutamatergic neurons (17.58% vGlut1-positive) **(Figure 3 A5)**. In line with the viral approach, OTR^+^ neurons constitute only a small population of 1.35% of all neurons (**Figure 2A**). Amongst the entire GABAergic and glutamatergic population, OTR^+^ neurons make up 6.82% and 0.16% respectively (**Figure 3 A6**).

**Figure 3.**
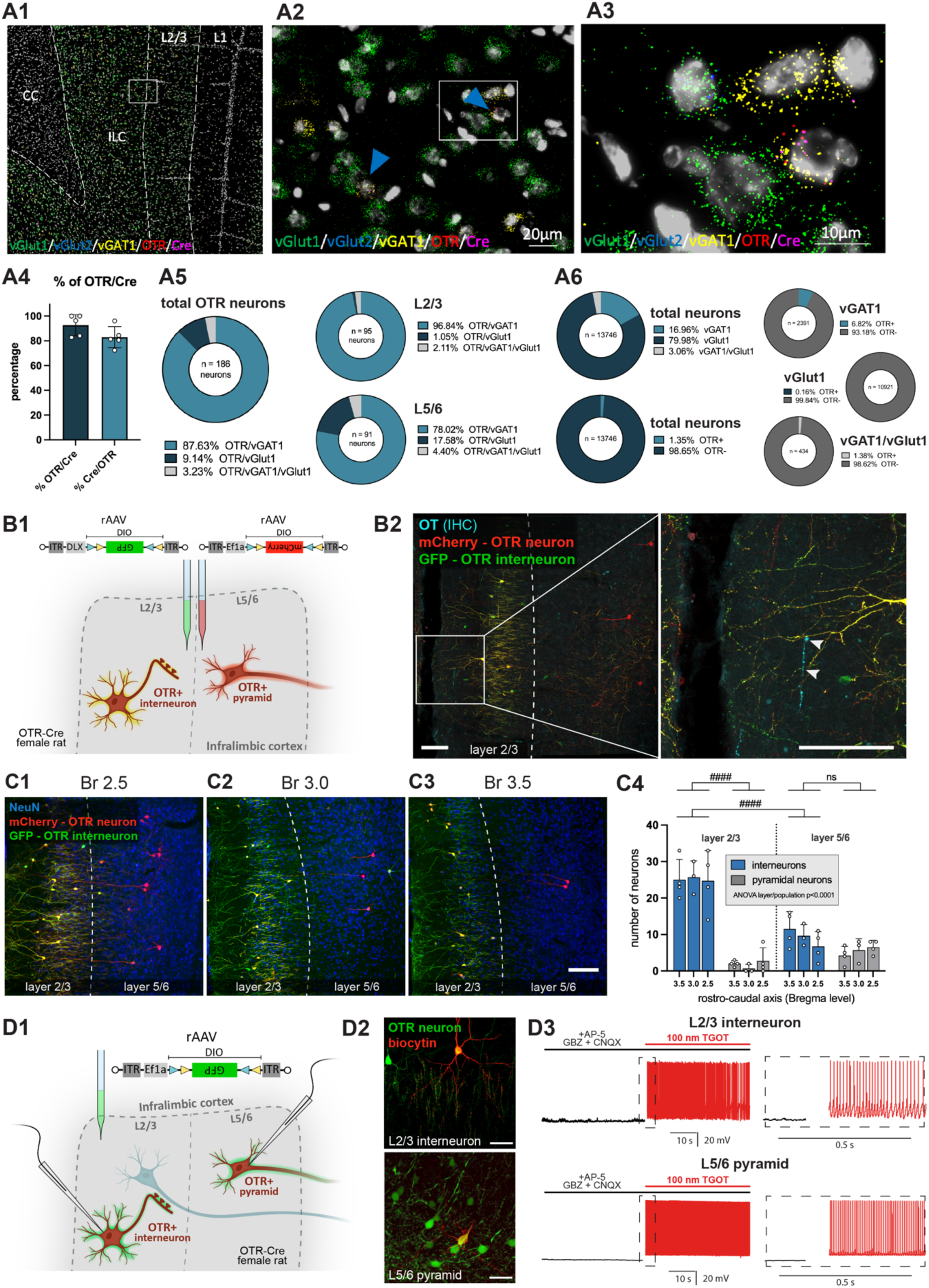
OTR^+^ cell types, their apposition to OT axons, and *ex vivo* responses to bath application of an OTR agonist. **A** RNAscope based characterization of OTR^+^ neurons in the ILC. **A1** Probes for vGlut1 (green), vGlut2 (blue), VGAT1 (yellow), OTR (red) and Cre (magenta) were used to visualize mRNA co-expression in the ILC of female OTR-Cre rats (n=5 animals). **A2-A3** Magnifications show representative vGAT1-positive neurons co-expressing Cre and OTR. **A4** Colocalization of OTR and Cre mRNA shows high specificity of the transgenic Cre rat line. **A5** The majority of OTR^+^ neurons in the ILC are GABAergic interneurons, with a small fraction of glutamatergic neurons, which are predominantly found in the layers 5/6. **A6** *In-situ-* hybridization shows that 1.35% of all neurons in the ILC are OTR^+^ neurons, with 6.82% of all GABAergic neurons being OTR^+^. **B** Viral vector-based characterization of OTR^+^ neurons in the ILC. **B1** Injection scheme depicting the injection of a mixture of two rAAVs: one expressing mCherry in a Cre-dependent manner to label the entire population of OTR^+^ ILC neurons and a second rAAV expressing GFP in a Cre-dependent manner only in OTR^+^ interneurons (utilizing the DLX-enhancer for specific expression in interneurons) into the ILC of OTR-Cre female rats. **B2** Representative confocal image depicting OTR^+^ interneurons dominating layers 2/3 (yellow), and individual OTR^+^ pyramidal neurons localized in layers 5/6 (red), with OT fibers (blue, white arrows) in both subdivisions of the ILC. Scalebar 100µm. **C** Quantification of OTR^+^ neurons in layer 2/3 and layer 5/6 along the rostro-caudal axis at **C1** Bregma 2.5, **C2** Bregma 3.0 and **C3** Bregma 3.5, Scalebar 100µm. **C4** Semi-automated, high-throughput analysis of the number of OTR^+^ neurons identified as interneurons (yellow) or pyramidal neurons (red) in the ILC (n=4 animals, each with 3 sections at three different bregma levels). **D** *Ex vivo* electrophysiological characterization of OTR^+^ neurons in the ILC. **D1** Injection scheme depicting the injection of a rAAV expressing GFP in a Cre-dependent manner in OTR-Cre animals for the identification of OTR^+^ neurons during *ex vivo* patch clamp recordings. **D2** Biocytin filled layer 2/3 interneuron and layer 5/6 pyramidal neuron. Scalebar 50µm. **D3** Representative traces of OTR^+^ neurons firing upon bath application of an OTR agonist (TGOT) (n=9 cells recorded from 4 rats for interneurons, n=5 cells recorded from 2 rats for pyramidal neurons). Statistical significance is indicated as * p<0.05, ** p<0.01, *** p<0.001, **** p<0.0001. Statistical significance of post-hoc tests is indicated as # p<0.05, ## p<0.01, ### p<0.001, #### p<0.0001. Error bars show standard deviation. For details on statistical tests please refer to Supplemental Document 1.

To complementarily characterize the distinct glutamatergic and GABAergic OTR^+^ populations in the ILC, we injected both a rAAV labelling all OTR^+^ neurons in red (rAAV-Ef1a-DIO-mCherry) as well as a second rAAV labelling all OTR^+^ GABAergic interneurons (specificity achieved by DLX-enhancer (Dimidschstein, Chen et al. 2016)) in green (rAAV-DLX-DIO-GFP, as all OTR^+^ interneurons will also be OTR^+^ neurons, they will appear yellow) into the ILC of female OTR-Cre rats **(Figure 3 B1)**. Similar to our findings with the RNAscope approach, we could identify an OTR^+^ interneuron population in layer 2/3 (yellow) and a mixed OTR^+^ pyramidal (red) and interneuron population in layer 5/6 **(Figure 3 B2, Extended Data Figure S3A)**. The results of semi-automated quantification of OTR^+^ neurons in distinct cortical layers (2/3 and 5/6) and along the rostro-caudal axis (Bregma 2.5, 3.0, 3.5), resembled the findings of in situ hybridization: in layer 2/3, 92.93% of OTR^+^ neurons are interneurons, while layer 5/6 also has a larger population of 37.04% pyramidal neurons (**Figure 3 C**). No significant differences of OTR^+^ neuron distribution were found along the rostro-caudal axis. Together, these experiments demonstrate that the largest population of OTR^+^ neurons in the ILC consists of GABAergic interneurons located in layer 2/3.

Histological quantification after viral labelling of OTR^+^ cell types confirmed that the majority of OTR^+^ neurons were GAD-expressing interneurons immunopositive for parvalbumin and/or calbindin **(Extended Data Figure S3 B)**. Detailed anatomical imaging of cell bodies of OTR^+^ neurons and OT fibers revealed the localization of OT-immunoreactive axons in close proximity to OTR^+^ neurons in different layers, pointing to axonal release as a probable source of OT acting on ILC OTR^+^ neurons and corroborating our aforementioned optogenetic experiments **(Extended Data Figure S3 C)**. Notably, OT fibers were found to be significantly closer to OTR^+^ neurons in layer 2/3 compared to layer 5/6 (0.04±0.07µm vs. 0.91±0.55µm, p<0.0001) **(Extended Data Figure S3 D)**.

The verify the presence of functional OTRs, we performed an *ex vivo* patch clamp experiment, where OTR^+^ neurons were virally labelled by rAAV-Ef1a-DIO-GFP **(Figure 3 D1)**, and classified based on their firing pattern, localization, and biocytin-based morphology visualization **(Figure 3 D2)**. All tested cells (interneurons n=9; pyramids n=5) fired extensively in response to bath application of selective OTR agonist (20-100nM TGOT) **(Figure 3 D3)**.

### Activity of ILC OTR^+^ interneurons promotes both social interaction and social preference

Given that the major population of ILC OTR-expressing neurons consists of inhibitory cells (primarily in layers 2/3), we aimed to investigate whether modulation of their activity induces similar behavioral effects as recruiting all ILC OTR^+^ neurons **(Figure 2C)**. To achieve this, we employed a rAAV equipped with the interneuron-specific enhancer DLX (Dimidschstein, Chen et al. 2016) to express Cre-dependent DREADDS (either hm3Dq(Gq) for activation or hM4Di(Gi) for inhibition) in OTR^+^ interneurons (rAAV-DLX-DIO-hm3Dq-GFP or rAAV-DLX-DIO-hm4Di-GFP) of the ILC **(Figure 4 A1)**. Post-hoc IHC analysis verified the main population of OTR^+^ interneurons being located in layer 2/3 **(Figure 4 A2)**. The two groups of rats (Gq: activation or Gi: inhibition) received CNO (3mg/kg) or saline i.p. injections 40 minutes before being introduced into a free social interaction session with an unknown conspecific. Chemogenetic activation of OTR^+^ interneurons significantly increased social interaction time (61.5±17.7s saline vs. 77.0±16.4s CNO, p=0.0346), while chemogenetic inhibition significantly decreased social interaction time (61.0±16.6s saline vs. 42.7±13.3s CNO, p=0.0197) **(Figure 4B)**. Investigation of a toy rat was not altered by either activation or inhibition of OTR^+^ interneurons (activation: 35.7±15.6s vs. 40.8±19.7s, p=0.4381; inhibition: 39.2±16.0s vs. 49.0±28.5s, p=0.4372). Given the previously indicated anxiolytic effect of OT action in OTR^+^ interneurons of male mice (Li, Nakajima et al. 2016), we performed OF and elevated zero maze (EZM) tests to investigate the putative effect of ILC OTR^+^ interneuron modulation on anxiety levels in female rats. Notably, chemogenetic activation of these cells changed neither anxiety nor locomotion parameters in our experiments **(Extended Data Figure S4 A-B)**. Moreover, we conducted a social novelty preference test and observed no difference induced by chemogenetic activation of ILC OTR^+^ interneurons (corner: p=0.0100, treatment: p=0.1474, corner x treatment: p=0.1937, post-hoc tests all ns) **(Extended Data Figure S4C)**. Taken together, these experiments showed that recruitment of ILC OTR^+^ interneurons is sufficient to promote social behavior without alterations of social novelty preference or anxiety.

**Figure 4.**
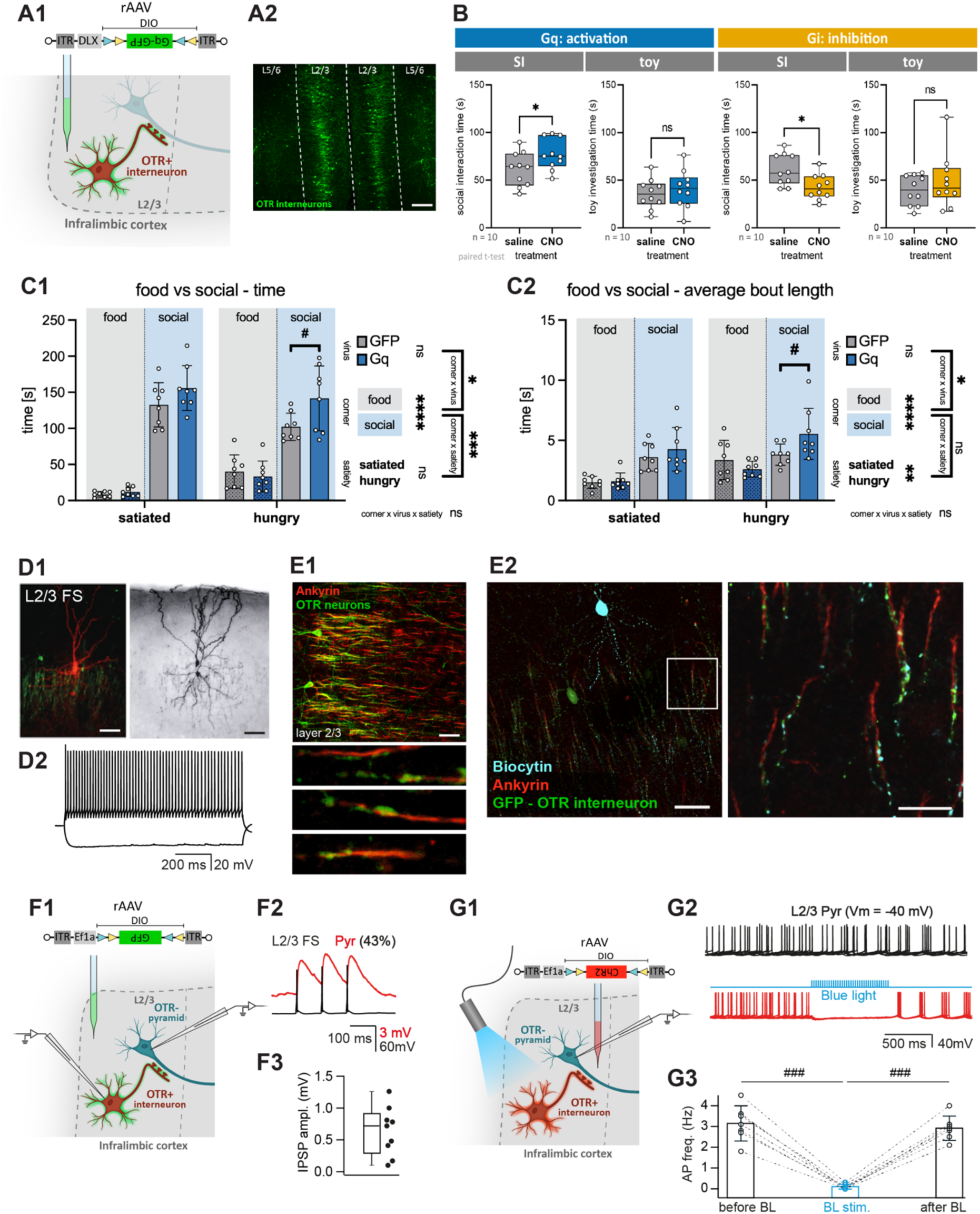
OTR^+^ interneurons modulate social behavior. **A** Chemogenetic manipulation of OTR^+^ interneurons in the ILC. **A1** Injection scheme depicting the injection of a rAAV expressing the excitatory (Gq) or inhibitory (Gi) DREADD (designer receptor exclusively activated by designer drugs) in a Cre-dependent manner in all OTR^+^ interneurons (via DLX enhancer) in the ILC of transgenic OTR-Cre female rats. **A2** Injection site showing localization of OTR^+^ interneurons (green) in layer 2/3. **B** Chemogenetic manipulation of OTR^+^ interneurons via CNO injection before a 5 min social interaction (SI) session with an unknown conspecific in the open field. Activation (n=10 animals) or inhibition (n=10 animals) of OTR interneurons significantly increases or decreases social interaction time, respectively. Investigation time of a toy rat in the open field remains unaltered by chemogenetic manipulation. **C** Food vs. social paradigm**. C1** Chemogenetic activation of OTR^+^ interneurons increases time spent investigating the social corner (n=8 animals). **C2** This effect is driven by an increase in the average length of bouts. **D** Morphological and functional characterization of OTR^+^ Chandelier neurons. **D1** Fluorescent (left) and DAB (right) staining of a biocytin-filled OTR^+^ ILC neuron reveal the morphological features of Chandelier cells bearing vertically aligned axonal cartridges. **D2** Representative *ex-vivo* recording of OTR^+^ Chandelier interneuron showing its fast-spiking (FS) activity. **E1** Ankyrin staining (red, marker for axon-initial segments) reveals the close proximity of OTR^+^ boutons (green, viral labelling) to the initial segment of the axon of the local OTR-negative (OTR^-^) pyramidal neurons. Scalebar 50µm. **E2** Biocytin filling (blue) of OTR^+^ neurons (green) and staining for ankyrin (red) verifies typical Chandelier morphology in 12 of 14 OTR^+^ interneurons. Scalebar 50µm overview, 25µm zoom. **F** Paired patch-clamp recordings of OTR^+^ fast-spiking (FS) interneurons. **F1** Injection scheme depicting the injection of a Cre-dependent GFP-expressing rAAV into the ILC of OTR-Cre rats to characterize OTR^+^ neurons; an *ex vivo* paired patch clamp experiment was performed, where OTR^+^ interneurons (green) were stimulated while the activity of OTR^-^ pyramidal neurons (no fluorescent marker) was recorded. **F2** A representative paired recording shows stimulation of the layer 2/3 FS OTR^+^ interneuron (black) and the evoked postsynaptic potential of an OTR^-^ pyramidal neuron (red). **F3** Post-synaptic potential (PSP) amplitude of OTR^-^ pyramids after stimulation of OTR^+^ interneurons (number of tested pairs: 21, number of connected pairs: n = 9, brain slices obtained from 3 rats). **G** *Ex vivo* optogenetic stimulation of OTR^+^ interneurons and patch clamp recording of OTR^-^ pyramidal neurons. **G1** Injection scheme depicting the injection of a Cre-dependent ChR2-expressing rAAV into the ILC of OTR-Cre rats, in order to activate OTR^+^ neurons upon BL stimulation; an *ex vivo* patch clamp experiment was performed, where OTR^+^ neurons (red) were stimulated with BL while the activity of OTR^-^ pyramidal neurons (no fluorescent marker) was recorded. **G2** Representative traces of a layer 2/3 OTR^-^ pyramidal neuron at baseline (black) and during BL stimulation of OTR^+^ neurons (red). **G3** Action potential frequency (AP freq.) of inhibited OTR^-^ pyramids before, during, and after optogenetic stimulation of OTR^+^ neurons (n = 7 cells, brain sections were obtained from 4 rats). Statistical significance is indicated as * p<0.05, ** p<0.01, *** p<0.001, **** p<0.0001. Statistical significance of post-hoc tests is indicated as # p<0.05, ## p<0.01, ### p<0.001, #### p<0.0001. Error bars show standard deviation. For details on statistical tests please refer to Supplemental Document 1.

To further elucidate the social motivational aspect of the observed behavior, we analyzed whether manipulation of the OTR^+^ interneuron population has an effect on social preference over another rewarding stimuli. Therefore, we subjected animals to an adapted social vs. food preference paradigm (Netser, Meyer et al. 2020) **(Extended Data Figure S4 E1-2)**, where in a satiated state animals typically prefer social interaction, while food restriction for 24 hours increases the interest in food. Rats were injected either with the chemogenetic excitatory rAAV system (rAAV-DLX-DIO-hm3Dq-GFP, ‘**Gq’**) previously used (Figure 4 A1), or with a rAAV lacking the Gq sequence as a control (rAAV-DLX-DIO-GFP, ‘**GFP’**). In the food vs. social preference experiment we observed that animals spent significantly more time at the social compared to the food corner (main effect corner, p<0.0001), and that food deprivation altered allocation across corners as expected (more time spent investigating the food corner after 24hrs of food restriction; satiety × corner, p=0.0004). Most importantly, that chemogenetic activation influenced the interaction time with the corners in a group specific manner (virus × corner, p=0.0431), pointing to a significant increase in social corner interaction time independently of satiety state (no interaction of virus x corner x satiety state p=0.2387). Additional post-hoc tests showed greater social corner time in Gq vs. GFP condition under deprivation (p=0.0155), with no group difference in food corner time (adjusted p=0.9766) (Figure 4 **C1**). Notably, this effect was driven by an increased average length of bouts (average length per visit of the social corner, virus × corner, p=0.0475) after CNO injection **(Figure 4 C2)**, while the number of bouts (number of visits to the social corner) **(Extended Data Figure S4 E3)** remained unaltered. To further investigate the effect of ILC OTR^+^ interneurons on non-social interest in a non-competing context we performed a similar test where only food was offered in one of the corners, with the second corner being empty. In this experiment, food deprivation had a significant effect on food investigation time (main effect food restriction, p=0.0448), but chemogenetic activation of OTR^+^ interneurons did not alter it (main effect virus, p=0.6628). The number of bouts of food investigation, as well as distance travelled and speed during the test were also not affected **(Extended Data Figure S4 D1-4)**. Together, these findings indicate that when a rat is presented with conflicting interests, ILC OTR⁺ interneurons promote social engagement in the hungry state, overriding competing homeostatic demands while not exerting a direct anorexigenic influence.

### OTR^+^ fast-spiking interneurons inhibit pyramidal neurons in the ILC

The aforementioned *ex vivo* patch clamp experiments and consecutive biocytin-based morphological analysis revealed that most layer2/3 OTR^+^ cells in the ILC display fast-spiking firing patterns (83%, n=24 patched cells, slices obtained from 5 rats) and exhibits axonal arborization characteristic for axo-axonic interneurons (Chandelier cells) (85.7%, n=14) (Somogyi 1977, Freund and Gulyás 1996) **(Figure 4 D)**. Bath application of TGOT at both low and high concentrations (20nM and 100nM, respectively) during *ex vivo* patch clamp recordings reliably and significantly increased their firing activity. Note that although the percentage of responding cells did not depend on the TGOT concentration, the average firing rate was twice as high at higher agonist concentrations **(Extended Data Figure S4F)**. Furthermore, IHC staining for ankyrin (a marker for the axon initial segment) revealed that the terminals of axon arbors of the OTR^+^ neurons form distinct arrays (so-called “cartridges”) along the axon initial segments of OTR-negative (OTR^-^) pyramids **(Figure 4E)**. Based on these morphological and functional features we could identify the vast majority of OTR^+^ interneurons in layer 2/3 of the ILC as predominantly Chandelier cells.

To functionally test whether OTR^+^ interneurons in layer 2/3 provide an inhibitory synaptic input to the neighboring pyramidal neurons, we performed *ex vivo* paired patch-clamp recordings in brain slices. Three weeks following an injection of a rAAV expressing GFP in a Cre-dependent manner into the ILC (rAAV-Ef1a-DIO-GFP), we searched for synaptic connections between fluorescently labelled OTR^+^ neurons and non-fluorescent OTR^-^ pyramidal cells in layer 2/3 **(Figure 4 F1-2)**. We observed that stimulation of putatively presynaptic OTR^+^ interneurons led to generation of a post-synaptic potential in 43% of recorded pairs with latency indicative of monosynaptic connections **(Figure 4 F3)**. Additionally, we applied a population-based *ex vivo* approach, utilizing optogenetic stimulation of ChR2-expressing OTR^+^ neurons (in rAAV-Ef1a-DIO-ChR2-mCherry ILC injected OTR-Cre rats) while simultaneously recording layer 2/3 OTR^-^ pyramidal neurons **(Figure 4 G1)**. Consistent with the paired-patch recordings, BL stimulation resulted in a significant (p<0.0001) decrease of action potential frequency in all recorded local output pyramidal neurons **(Figure 4 G2-3)**. In a follow-up *ex vivo* patch clamp experiment we recorded spontaneous excitatory (EPSC) and inhibitory (IPSC) post-synaptic currents in layer 2/3 OTR^-^ pyramidal neurons. The bath application of TGOT significantly increased the IPSC amplitude, with no effect on EPSC amplitude or EPSC and IPSC frequency **(Extended Data Figure S4G)**, further corroborating the inhibitory nature of the synaptic connection of OTR^+^ neurons in layer 2/3 targeting OTR-negative pyramidal neurons. Together, these experiments suggest that endogenous OT release in the ILC activate layer 2/3 OTR^+^ interneurons, which inhibit neighboring OTR^-^ down-streaming principal cells.

### OTR^+^ interneurons inhibit the ILC−basolateral amygdala pathway

Having established the targeted inhibitory input from OTR^+^ interneurons to layer 2/3 OTR^-^ pyramidal neurons, our next objective was to investigate their potential downstream projection sites. Therefore, we analyzed the neuronal activity marker cFos expression in seven pre-selected brain regions downstream to the ILC (Gabbott, Warner et al. 2005), which were also reported to be functionally involved in modulation of social behaviors (Ko 2017), upon the chemogenetic activation (rAAV-DLX-DIO-Gq-GFP) of ILC OTR^+^ interneurons. While control animals receiving saline injections showed no specific cFos expression in OTR^+^ interneurons (0.69±0.37 % cFos/GFP double-positive cells, n=375 GFP^+^ neurons from 3 animals), animals receiving CNO showed cFos signal in these GFP+ cells (99.91±0.15 % cFos/GFP double-positive cells, n=424 GFP^+^ neurons from 3 animals) **(Figure 5 A1)**. To allow for unbiased quantification of neuronal cFos expression, we employed a fully automated pipeline based on the Imaris software which analyzed cFos signals in more than 500,000 neurons across the seven different brain regions. Activating OTR^+^ inhibitory interneurons in the ILC resulted in an overall decrease of cFos expression in the ILC (main treatment effect, p=0.0004), with a significant decrease in layer 2/3 (post-hoc layer 2/3, p<0.0001, post-hoc layer 5/6 0.0604) **(Figure 5 A2)**, indicating extensive inhibition of OTR(GFP)-negative principal cells **(Figure 4G)**, particularly in layer 2/3 where the main population of OTR^+^ interneurons resides. As for the analyzed downstream regions, cFos expression was not significantly changed in the periaqueductal grey (PAG), lateral septum, supramammillary nuclei (Summ), or ventral tegmental area (VTA). However, it was significantly reduced in the basolateral amygdala (BLA) (15.4±4.9% vs. 8.7±2.0%; p=0.0185) **(Figure 5 A3)**. This suggests that experimental OT release in the ILC ultimately reduces the activity of BLA neurons. Notably, in the same paradigm, we observed a significant increase in cFos expression in the nucleus accumbens (NAc) (10.7±2.3% vs. 22.2±7.9%; p=0.0084), hinting at differences between the possible functional dichotomy of BLA and NAc projections from the ILC. This may suggest that ILC output neurons projecting to the NAc are not directly (monosynaptically) inhibited by layer 2/3 OTR^+^ interneurons.

**Figure 5.**
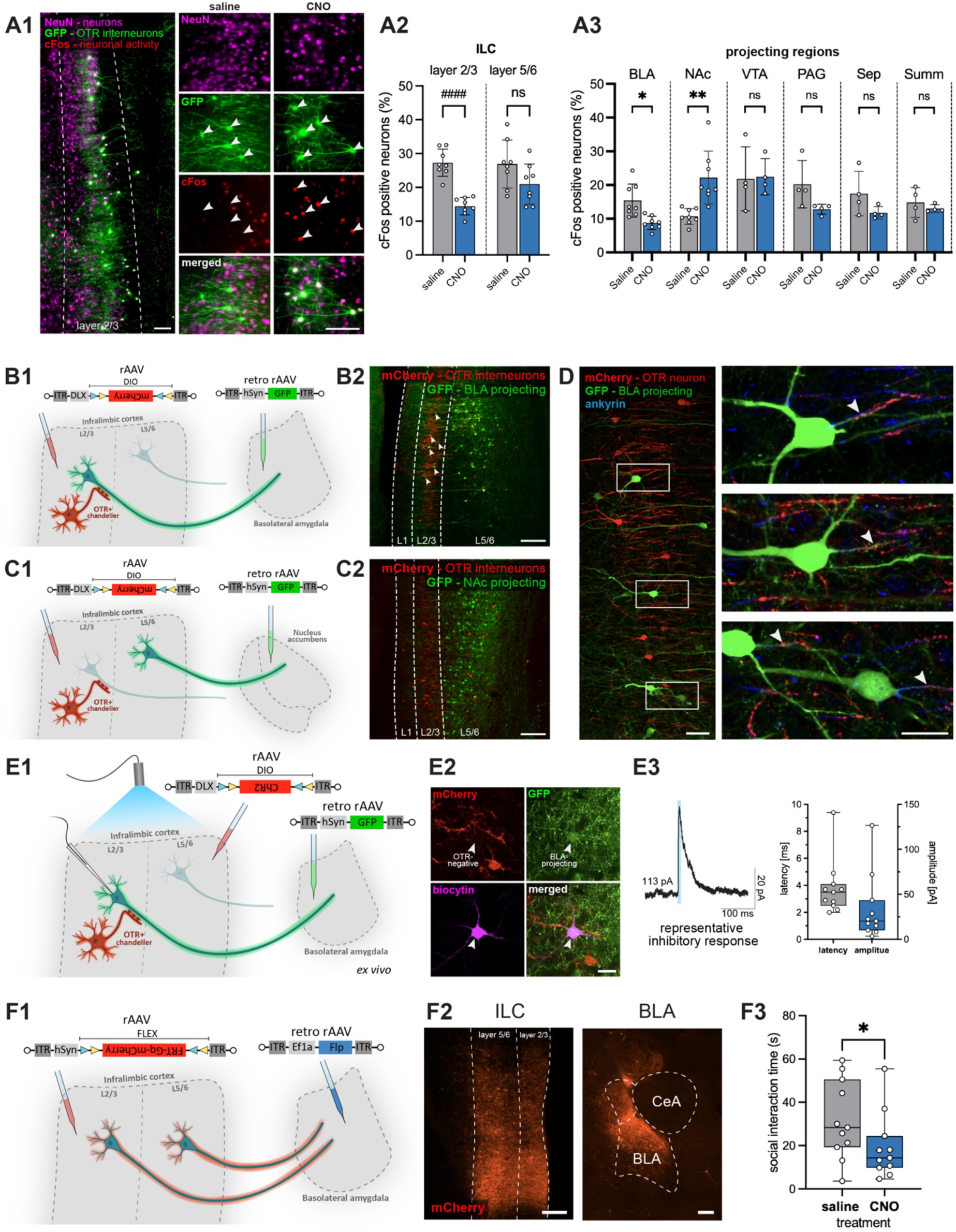
OTR^+^ interneurons modulate neuronal activity in the BLA. **A** Expression of cFos after chemogenetic activation of OTR^+^ interneurons in the ILC. **A1** A rAAV expressing Gq-GFP (excitatory DREADD) in all OTR^+^ interneurons (green) was injected into the ILC of female rats; after either CNO or saline administration followed by perfusion (100 minutes post injection) brain sections were immunohistochemically stained for NeuN (magenta) and cFos (as a marker for neuronal activity, red). After CNO injection, OTR^+^ interneurons were activated chemogenetically as verified by cFos expression (white arrows), whereas after saline injection only baseline cFos expression occurred with no cFos expression in OTR^+^ interneurons (white arrows). Scalebar 100µm **A2** Local reduction of cFos expression in the ILC after chemogenetic activation of OTR^+^ interneurons in the ILC, specific to layer 2/3. **A3** Local reduction or increase of cFos expression in the BLA or NAc, respectively, after chemogenetic activation of OTR^+^ interneurons in the ILC. No significant change of cFos expression was observed in the VTA, PAG, septum, or Summ (brain sections were obtained from n=8 animals per group). **B** Retrograde tracing of BLA-projecting neurons in the ILC. **B1** Injection scheme depicting the injection of a retrogradely GFP-expressing rAAV into the BLA with simultaneous injection of a Cre-dependent mCherry-expressing interneuron-specific rAAV into the ILC. **B2** BLA-projecting neurons are found both within layer 5/6, as well as within in the “inhibitory sleeve” of OTR^+^ interneurons in layer 2/3 (white arrows). Scalebar 200µm. **C** Retrograde tracing of NAc-projecting neurons in the ILC. **C1** Injection scheme depicting the injection of a retrogradely GFP-expressing rAAV into the NAc with simultaneous injection of a Cre-dependent mCherry-expressing interneuron-specific rAAV into the ILC. **C2** NAc-projecting neurons are found only within layer 5/6, but not within the “inhibitory sleeve” of layer 2/3. Scalebar 200µm. **D** Representative axo-axonic contacts (white arrows) of OTR^+^ interneurons (red) on BLA-projecting pyramidal neurons (green). The axon-initial segment is stained immunohistochemically for ankyrin (blue). Scalebar 50µm overview, 25µm zoom. **E** Electrophysiological patch-clamp recording of OTR^-^ neurons projecting to the BLA. **E1** Injection scheme depicting the injection of a rAAV expressing ChR2 in a Cre-dependent manner in OTR^+^ interneurons of OTR-Cre animals, with additional injection of a rAAV into the BLA for retrogradely identifying projecting neurons. During an *ex vivo* patch clamp recording of BLA-projecting neurons, BL is used to activate OTR^+^ interneurons. **E2** Representative image of a biocytin-filled OTR^-^ BLA-projecting pyramidal neuron. Scalebar 20µm. **E3** Representative averaged trace of a whole-cell patch clamp recording (voltage clamp, holding voltage -60mV) depicting an inhibitory response of a BLA-projecting neuron within the “inhibitory sleeve” (layer 2/3) after optogenetic activation of OTR^+^ interneurons in the ILC. Latency and amplitude of recordings for all neurons (n=11 recorded neurons). **F** Social interaction after chemogenetic activation of BLA-projecting neurons originating in the ILC. **F1** Injection scheme depicting the injection of a retrogradely Flp-expressing rAAV into the BLA with simultaneous injection of a Flp-dependent (Frt) Gq-mCherry-expressing rAAV into the ILC to chemogenetically activate BLA-projecting ILC neurons after i.p. application of CNO. **F2** Representative injection site showing expression of the rAAV-hSyn-FRT-Gq-mCherry in cell bodies and fibers in the ILC and projecting fibers in the BLA. Scalebar 250µm. **F3** Chemogenetic manipulation of BLA-projecting neurons via CNO injection before a 5 min social interaction (SI) session with an unknown conspecific in the open field. Activation of BLA-projecting neurons decreases social interaction time (n=11 animals). Statistical significance is indicated as * p<0.05, ** p<0.01, *** p<0.001, **** p<0.0001. Statistical significance of post-hoc tests is indicated as # p<0.05, ## p<0.01, ### p<0.001, #### p<0.0001. Error bars show standard deviation. For details on statistical tests please refer to Supplemental Document 1.

To test this hypothesis, we performed viral retrograde tracing of neurons projecting to the BLA or to the NAc by injection of a retrograde GFP-expressing rAAV (retro-rAAV-hSyn-GFP) into either BLA or NAc. The same animals were injected into the ILC with a Cre-dependent rAAV expressing mCherry in OTR^+^ interneurons (rAAV-DLX-DIO-mCherry) to visualize the overall topography of back-labeled projecting cells in the ILC in relation to the OTR^+^ cell population **(Figure 5 B1, C1)**. Injection sites in the BLA or NAc were marked by co-injection of a small amount of soluble fluorescent beads in the tip of the injection pipette **(Extended Data Figure S5 A)**. A substantial number of BLA-projecting neurons was found within the “inhibitory sleeve” formed by the vertical axonal cartridges of OTR^+^ Chandelier cells in layer 2/3 (18.1±7.4 neurons per hemisphere in 50µm sections) as well as in the deeper layers of the ILC (31.8±12.1 neurons per hemisphere in 50µm sections) **(Figure 5 B2, Extended Data Figure S5 C)**. In contrast, retrogradely labeled NAc-projecting neurons were found exclusively in the deeper layers of the ILC, but not within the “inhibitory sleeve” of layer 2/3 **(Figure 5 C2)**. Furthermore, detailed morphological analysis of BLA-projecting neurons revealed that OTR^+^ interneurons form direct axo-axonic contacts with BLA-projecting pyramidal neurons in layer 2/3 (**Figure 5D, Extended Data Figure S5 B)**, but not with NAc-projecting pyramidal neurons in layer 5/6. Thus, we showed anatomical evidence of a direct and local OTR^+^-neuron-mediated inhibition of BLA-projecting neurons in layer 2/3 of the ILC, and focused on investigating this ILC-BLA downstream pathway.

Following this clear anatomical distinction, we next verified inhibitory action of ILC OTR^+^ interneurons on OTR^-^ pyramidal neurons projecting to the BLA. In the first set of experiments, we injected a retrograde rAAV expressing GFP into the BLA (retro-rAAV-hSyn-GFP) and, in parallel, a Cre-dependent rAAV expressing ChR2 in interneurons (rAAV-DLX-DIO-ChR2-mCherry) into the ILC of OTR-Cre-rats. After a 3-week expression period, OTR^+^ interneurons were optogenetically activated by BL stimulation in an *ex vivo* experimental setup, while BLA-projecting OTR^-^ pyramids were recorded in whole cell mode **(Figure 5 E1)**. Neurons were patched only if they were both GFP-positive (BLA-projecting) and mCherry-negative (OTR^-^ neurons) **(Figure 5 E2)**. Recorded neurons were identified as principal cells based on their typical discharge pattern and morphology after post-recording staining procedure **(Extended Data Figure S5 D and Figure 5 E2)**. Notably, all recorded neurons (n=11) responded with inhibition to BL induced OTR^+^ interneuron activation (**Figure 5 E3** for responses of BLA-projecting OTR^-^ pyramids, **Extended Data Figure S5 E** for representative trace verifying activation of OTR^+^ interneuron by BL).

Finally, we chemogenetically targeted the BLA-projecting population in the ILC, in order to verify its involvement in social behavior. We injected a rAAV expressing the excitatory DREADD Gq-mCherry in a Flp-dependent manner (rAAV-hSyn-FRT-Gq-mCherry) into the ILC of female rats, while simultaneously injecting a retro-rAAV expressing Flp (retro-rAAV-Ef1a-Flp) into the BLA (Figure 5 **F1**). With post-hoc verification of injection sites we could observe cell bodies in the ILC as well as dense fibers in the BLA (Figure 5 **F2**). Animals that received CNO injection prior to a free social interaction session spent significantly less time investigating the social partner (31.9±18.1s vs. 19.3±15.0s; p=0.042) (Figure 5 **F3, Extended Data Figure S5 F**). Thus, we could show that activating BLA-projecting neurons had the same effect on social interaction as inhibiting OTR^+^ interneurons in the ILC (Figure 4 **B,** right), supporting our hypothesis of OTR^+^ interneuron mediated inhibition of BLA-projecting neurons driving sociability in female rats.

## DISCUSSION

Our study demonstrates that OT release within the mPFC directly facilitates social behavior in female rats. We found that the ventral part of the mPFC (ILC) is preferentially innervated by OT fibers and enriched in OTR-expressing neurons, rendering it a major site of OT action. Intriguingly, we observed OT neuron axons across all cortical layers with the highest density in the layer 5/6, while the majority of OTR^+^ neurons resides in the layer 2/3. This apparent mismatch can be explained by the anatomical positioning of the ILC adjacent to the forceps minor of the rat corpus callosum, and long-range axons coursing within callosal/white-matter tracts that frequently send collaterals towards deep cortical layers (Heidbreder and Groenewegen 2003). Consistent with this architecture, OT projections arising from hypothalamic nuclei travel through major forebrain fiber bundles and elaborate fine collaterals in deep layers of the mPFC, producing higher apparent OT-fiber density in layers 5/6 than in superficial layers. This difference is especially apparent when using virus-based reporters, which reveal thin-caliber OT axons and terminals that are often below the detection threshold of classical peptide immunohistochemistry. The OT innervation detected in layer 1 aligns with the presence of dendritic arbors of layer 2/3 OTR^+^ Chandelier cells extending into the superficial cortex, a site of long-range modulatory inputs and distal dendritic integration (Inan and Anderson 2014). Given the capacity of OT to act via local diffusion at very low concentrations, OT released from sparse layer-1 afferents could effectively engage OTR^+^ interneurons in layers 2/3 despite the lower local fiber density (Chini, Verhage and Grinevich 2017). Notably, utilizing the novel GRAB OT-sensor (Qian, Wang et al. 2023) *ex vivo*, we showed OT binding upon optogenetic stimulation of OT fibers in the ILC, demonstrating local axonal OT release in this brain region.

*In vivo*, a similar optogenetic stimulation of OT fibers in the ILC increased social interactions between female rats. These findings are consistent with the involvement of OT in the modulation of social behavior through its actions in various brain regions (Marlin and Froemke 2016, Grinevich and Stoop 2018). Moreover, our results extend the recently reported analgesic OT input to the mPFC (Liu, Li et al. 2023), revealing its role in facilitating social interactions. Taken together with the prosocial effects of hypothalamic OT neuron stimulation (Lee, Rhim et al. 2016, Tang, Benusiglio et al. 2020), this work supports the role of OT in promoting social motivation and interaction.

We showed that OTR^+^ neurons in the ILC are intrinsically active during social behavior, with activity specific to interaction events initiated by the test rat, not the social partner (e.g., sniffing vs. being sniffed). This suggest their role in driving social interactions and may further indicate that ILC OTR^+^ neurons are crucial in the decision making processes underlying social executive functions of the mPFC (Kingsbury, Huang et al. 2019). Population imaging in the mouse mPFC shows that ensembles preferentially encode social olfactory cues and refine with experience (Levy, Tamir et al. 2019). This is consistent with a role for ILC circuits in driving social actions rather than merely reflecting sensory processing. It is tempting to propose a pivotal role of OT signaling in this process, yet a direct link between acute OT release and the ILC OTR⁺ calcium activity requires further targeted testing. Nevertheless, we proved the necessity of ILC OTR signaling for social interaction by local depletion of OTR. Our *in vivo* CRISPR/Cas9-mediated OTR targeting should be interpreted as a partial loss-of-function rather than a complete receptor knockout. First, viral transduction and CRISPR editing are inherently mosaic, so a fraction of OTR-expressing neurons likely retain intact receptors. Second, indel heterogeneity may generate alleles with partial activity. Third, residual receptor functionality unrelated to high-affinity ligand binding (for example heteromerization/dimerization) may persist and maintain some downstream signaling (Boender, Boon et al. 2023). These features likely explain the residual responsiveness to TGOT we detected (n=1 neuron, **Extended Data Figure S2 F**) and indicate that our behavioral effects probably underestimate the full impact of abolishing OTR signaling in the ILC. Our results demonstrate that OTR^+^ neuron activation in the ILC facilitates sociability via axonal OT release (Knobloch, Charlet et al. 2012).

We further explored the local ILC neuronal circuit receptive to OT using multiplex fluorescent *in situ* hybridization and neuroanatomical analyses. Our findings revealed that OTR expression in the ILC is restricted to a small subset of neurons, underscoring their significant role in social behavior. This finding is in line with our previous work in the PVN, where we showed that a small population of parvocellular OT neurons modulates inflammatory pain processing (Eliava, Melchior et al. 2016). Taken together, this highlights the potent influence of OT in central circuits, even under conditions of low concentration or limited OTR expression. Additionally, we found that the vast majority of OTR^+^ neurons in the ILC are GABAergic interneurons, located predominantly in layers 2/3. This aligns with mouse mPFC reports emphasizing a predominantly interneuronal OTR population (Nakajima, Görlich and Heintz 2014, Li, Nakajima et al. 2016). By contrast, besides the interneuron population, Tan, Singhal et al. (2019) identified a substantial VGluT1⁺ OTR^+^ population in the mouse mPFC. We suspect these differences reflect a combination of species (rat vs. mouse), subregion (ILC vs. broader PFC/PL), sex (female rats), and methodological factors (driver lines, reporter/probe sensitivity, and positivity thresholds). Notably, our OTR-Cre specificity was validated by RNAscope (Cre↔OTR mRNA overlap ∼83-93%) and pharmacology (all OTR^+^ neurons marked with GFP responded to bath application of TGOT in our patch clamp experiments). Based on morphological and electrophysiological features, we found that the majority of these OTR^+^ neurons are Chandelier cells. These GABAergic interneurons are powerful inhibitors of output principal cells in cortical circuits (Jung, Choi and Kwon 2022). They form unique axo-axonic synapses, quenching the generation of action potentials at the axon initial segment of pyramidal neurons. This unique inhibitory efficacy of Chandelier cells enables their precise control over the activity of output pyramidal neurons and overall network excitability. The functional activation of these interneurons in our experiments led to significant behavioral changes, suggesting that GABAergic modulation within the ILC is crucial for social behavior regulation. This finding is consistent with the known role of GABAergic interneurons in shaping cortical output and influencing behavior (Yang, Mack et al. 2021). Notably, we did not observe any effects on anxiety behaviors when activating ILC OTR^+^ interneurons. This is in line with work from Li, Nakajima et al. (2016), who showed that activating OTR-expressing interneurons in the mPFC reduces anxiety-like behavior in males but not females (with OTR loss producing the opposite, male-specific anxiogenic effect). In contrast, the same manipulation enhances social behavior in females but not males, revealing complementary sex-specific routing of OT’s influence on anxiety vs. sociability, a sex difference yet unexplored in rats.

Considering that the activation of ILC OTR⁺ interneurons increased investigation of the social, but not the food corner in the social vs. food paradigm, the observed change can be interpreted as facilitation of social approach under competing motivation rather than an anorexigenic effect. This choice-specific engagement supports a selective action of OTR^+^ neuron modulation, rather than a global change in arousal or locomotion. The observed shift towards increased social investigation even in the presence of food after 24hrs of food deprivation is consistent with the hypothesis of mutually inhibitory ensembles in the mPFC (Gangopadhyay, Chawla et al. 2021). In line with their hypothesis and based on our results, we suggest that social decision making is facilitated by specific neuronal ensembles within the ILC that are selectively activated by social stimuli and modulated by OT action. Thus, specific OTR^+^ interneurons within the ILC promote social behavior and contribute to the encoding of socially relevant information and biasing decisions toward prosocial behaviors. In contrast, certain non-social ILC ensembles are activated by non-social stimuli and are involved in processing individual-centric information. Independent of OT signaling, work in rats shows that inactivation of the ILC broadly impaired both active and inhibitory avoidance and suppressed inappropriate reward seeking (Capuzzo and Floresco 2020). Consistent with this, a comprehensive review concluded that ILC function is highly context-dependent, varying with task demands and the specific neuronal ensembles recruited (Nett and LaLumiere 2021). In our social vs. food choice paradigm, activation of OTR⁺ ILC interneurons selectively increased social investigation without generalized arousal, consistent with the proposed role of the ILC in resolving competing motivations by suppressing non-social drives when social cues are behaviorally salient. Intriguingly, the prosocial effect of ILC OTR^+^ interneuron modulation emerges stronger under hunger. This suggests that OT signaling in the ILC acts not necessarily as a tonic enhancer of sociability, but as a context-dependent biasing mechanism that promotes social engagement specifically when motivational conflict is strong.

We also investigated subcortical regions downstream to OTR^+^ interneurons focusing on the BLA and the NAc, two areas that are well-known to be involved in socially relevant functions including fear, reward, and social interaction (Gabbott, Dickie et al. 1997, Gabbott, Warner et al. 2005, Bloodgood, Sugam et al. 2018). Chemogenetic activation of OTR^+^ ILC neurons in our experiment led to downregulation of cFos expression in the BLA and increased cFos expression in the NAc. We propose that the observed reduction in the number of cFos-positive cells in the BLA is caused by the ILC OTR^+^ interneurons-mediated inhibition of ILC principal cells providing excitatory input to the BLA. Among the mPFC subdivisions, the ILC in particular is known to target a dense population of GABAergic interneurons in the BLA (Davis, Zaki et al. 2017) and the BLA in turn densely innervates the NAc (Correia, McGrath et al. 2016). We hypothesize that the observed cFos upregulation in the NAc might be caused by the suppression of BLA interneuron activity, thereby increasing the excitatory signaling from the BLA to the NAc. As mapping circuits via cFos activity patterns comes with the caveat of spatial and temporal ambiguity, we further investigated the OTR-ILC-BLA/NAc circuit both anatomically and electrophysiologically. Retrograde tracing following injections in the BLA and NAc revealed BLA-projecting principal cells in the ILC that were embedded within a dense “inhibitory sleeve” and NAc-projecting principal cells only in deeper cortical layers. Additionally, we showed OTR^+^ interneurons forming axonal cartridges at the initial axonal segment of neurons projecting to the BLA which was not observed in the case of NAc-projecting principal cells. This was further confirmed by *ex vivo* studies which showed direct monosynaptic coupling between OTR^+^ interneurons and BLA-projecting principal cells. Notably, a study by Lu, Tucciarone et al. (2017) demonstrated that a subset of layer 2 Chandelier cells in the PrL cortex selectively innervates BLA-projecting principal cells (_BLA_PCs) over contralateral-cortex-projecting principal cells (_CC_PCs), while receiving preferential input from _CC_PCs, thereby establishing a directional inhibitory module that gates BLA-related output. Interestingly, they also found _BLA_PCs in the same laminar layers as Chandelier cells, while _CC_PCs were located in deeper layers, resembling our finding of BLA-vs. NAc-projecting neurons. This spatial coincidence suggests that OTR⁺ interneurons are well-positioned to modulate _BLA_PCs ensembles. While their study did not indicate a behavioral role of this neuronal subset, our work suggests its prosocial role. While unlikely, considering the absence of connections to NAc-projecting neurons we found in this study, we cannot exclude the possibility of OTR^+^ Chandelier cells projecting onto neurons in deeper layers. It has been reported previously that Chandelier cells in layer 2/3 innervate pyramidal neurons in cortical layers 2, 3, and 5 (Jiang, Wang et al. 2013, Lee, Wang et al. 2015), and that while they have mostly vertically oriented axonal segments in layer 2/3, individual long descending axons reached and arborized in layer 6 (Woodruff, Anderson and Yuste 2010). However, in our experiments, chemogenetic activation of ILC OTR^+^ interneurons did not alter cFos expression in deep ILC layers, suggesting that their effects are largely confined to layer 2/3.

Importantly, several studies indicate the ILC as a key node for social approach via amygdala-directed outputs. In mice, BLA-projecting ILC neurons are preferentially recruited by social cues, and normal sociability requires intact ILC-BLA signaling (Huang, Zucca et al. 2020). In hamsters, chemogenetic activation of the ILC-BLA connection during social defeat confers resilience and reduces defeat-associated behavior while lowering BLA c-Fos (Dulka, Bagatelas et al. 2020). Our finding that activating OTR^+^ interneurons in the rat ILC increases sociability, coupled with reduced BLA cFos expression, aligns with a model in which ILC microcircuits gate amygdala-directed output to favor social approach under challenge. This is further strengthened by the antisocial effects of stimulation of ILC-BLA-projecting pyramidal neurons we observed.

We acknowledge the possibility that, beyond the ILC-BLA circuit described here, other OT-dependent pathways originating in the mPFC may underlie distinct aspects of behavior. There are reports that principal neurons in the mPFC are sensitive to bath application of OT (Ninan 2011) or express OTRs (Tan, Singhal et al. 2019 and this study). In mice, numerous OT-sensitive principal cells can be found throughout the entire mPFC, including the infralimbic area (Newmaster, Nolan et al. 2020), and they are located in both superficial and deeper layers. On the contrary, our results in rats report considerably fewer (9.14% in situ hybridization data, Figure 3A) principal cells that express OTRs and these cells are located almost exclusively in the deeper layers, particularly outside of the layer 2/3 “inhibitory sleeve”. This may explain why we did not observe social novelty preference alterations following the activation of OTR^+^ interneurons **(Extended Data Figure S4C)** as previously found in male mice after manipulation of the direct mPFC-BLA pathway (Tan, Singhal et al. 2019). In addition to the differences in species and sex, Tan and colleagues showed that these principal OTR^+^ neurons specifically alter social memory, while having no particular effect on general sociability. This could indicate the existence of two independent, and potentially species-and sex-dependent OT-modulated pathways for sociability (interneurons) and social memory (pyramidal neurons). A distinctive organization of the OT systems in different mammals, including mice and rats, has been identified at the anatomical and behavioral level (Insel, Gingrich and Young 2001, Freeman and Young 2016, Althammer, Jirikowski and Grinevich 2018), reflecting species-dependent processing of social information and subsequent selection of optimal social strategy (Young 1999, Bielsky and Young 2004, Anacker and Beery 2013, Burkett, Andari et al. 2016).

In conclusion, our study identifies a critical role for OT signaling within the ILC in facilitating social behaviors in female rats **(Extended Data Figure S5 G)**. Activation of GABAergic OTR^+^ interneurons in particular appears to play a significant role in modulating social interaction. These findings contribute to our understanding of the neural mechanisms underlying social behavior and highlight the potential of targeting OT pathways for therapeutic interventions in social deficits (Meyer-Lindenberg, Domes et al. 2011), as the PVN/SON-to-mPFC OT signaling pathway is conserved throughout evolution and exists in several primate species including humans (Rogers, Ross et al. 2018, Lefevre, Meza and Miller 2024).

## Acknowledgments

This study was inspired by Professor Peter H. Seeburg (1944–2016), who encouraged us to investigate oxytocin signaling in cortical areas. Although its completion was delayed, we hope that our work fulfills his vision. We thank Andreas Draguhn for his input on electrophysiological experiments, Laura Dötsch for performing the Western blot experiment, Selina Wunsch for assisting in the cFos quantification, and Quirin Krabichler for technical assistance with virus injections. This work was supported by a PhD scholarship of the Studienstiftung des Deutschen Volkes to SS, Humboldt Research Fellowship to AK, Marie Sklodowska-Curie fellowship (101018877) to AL, IKY Scholarship, “Maria Zaousi Memorial” bequest, for postdoctoral research abroad in psychiatry to ARa, DFG Emmy Noether starting grant AL 2466/2-1 to FA, the Synergy European Research Council (ERC) grant “OxytocINspace” 101071777 to VG, SFB Consortium 1158-3 to VG, DFG grant 533293533 to VG, and German-Israeli Project cooperation (DIP) GR3619-1 to RH, VG and SW, and ERANET-Neuron grant GR 3619/25-1 to VG and SW.

## Authors contribution

Project conception: ME, VGrin. Anatomical experiments: ME, SS, FA, AK, KA, TS, FS, YP. Behavioral experiments (optogenetic/chemogenetic): SS, AK, AL, ARa, KA, AZ, HS. *Ex vivo* patch-clamp electrophysiology: AK, JL, ARo, KA, AC, DH, SM, HM. *Ex vivo* oxytocin sensor: AC, CD, HP, VGrel, K-YW, AC. *In vivo* fiber photometry: AL, RP, AS. RNAscope: AK, AG, AT, KA, ABla. Methodology: SS, AK, JS, ABoe, AL, RH, RT, SN, SW, LG, YL, AC. Manuscript preparation: SS, AK, VGrin. Project administration and supervision: ME and VGrin. Funding acquisition: VGrin.

## METHODS

### Animals

Female Sprague–Dawley rats (3–8 weeks, Janvier Labs) and transgenic OTR-Cre rats bred in-house (Central Institute for Mental Health, Mannheim) were used. Adult virgin females (8–16 weeks) were included in anatomical, functional, and behavioral experiments. For single-day behavioral tests (free social interaction, between-subject design), only animals in diestrus were included, as described previously in Tang, Benusiglio et al. (2020). Rats were maintained under a 12-h light/dark cycle (lights on 07:00), 22–24 °C, 50 ± 5% humidity, group-housed in EU Makrolon type IV cages, with food and water ad libitum unless stated differently. Rats were group-housed under a 12-h light/dark cycle (lights on 07:00, 22–24 °C, 50 ± 5% humidity) with food and water ad libitum unless stated otherwise. All procedures complied with German legislation and were approved by the Baden-Württemberg Animal Ethics Committee (licenses G-193/20 and G-55/23).

#### Transgenic OTR -Cre Rats

Transgenic OTR-Cre rats (Iwasaki, Lefevre et al. 2023) express Cre recombinase in all OTR-expressing neurons, enabling cell type–specific expression of Cre-dependent (double-floxed inverted orientation, DIO) AAVs. Animals were generated by breeding heterozygous Cre⁺ males with wild-type females, and offspring were genotyped using a PCR-based protocol to identify heterozygous Cre⁺ or homozygous Cre⁻ animals.

### Viral Vectors

Recombinant adeno-associated viruses (rAAVs) of serotype 1/2 were used following previously established methodologies (Knobloch, Charlet et al. 2012). HEK293T cells for viral production were obtained from Addgene (catalog number 240073). The OT-promoter controlled rAAV-OTp-ChR2-mCherry, rAAV-OTp-Venus, and rAAV-OTp-ChrimsonR-mScarlet, as well as the Cre-dependent rAAV-Ef1a-DIO-GFP, rAAV-Ef1a-DIO-mCherry, rAAV-Ef1a-DIO-GCaMP6s, and rAAV-Ef1a-DIO-ChR2-mCherry, were produced in-house (titers between 10⁹ and 10¹⁰ genomic copies/μl). The plasmid for rAAV-hSyn-FLEX-FRT-Gq-mCherry was obtained from Addgene and the viruses were produced in-house (titer > 10⁹ genomic copies/μl). rAAVs for interneuron targeting (rAAV-DLX-DIO-GFP, rAAV-DLX-DIO-mCherry, rAAV-DLX-DIO-Gq-GFP, rAAV-DLX-DIO-Gi-GFP, rAAV-DLX-DIO-ChR2-mCherry) were purchased from the ETH Zurich Viral Vector Facility (serotype DJ, titers 6.5 × 10¹² – 8 × 10¹² vg/ml). The serotype 9 rAAVs used for OTR knockdown (rAAV-hSyn-spCas9, rAAV-u6-gRNA(OTR)-hSyn-mCherry, rAAV-u6-gRNA(lacZ)-hSyn-mCherry) were obtained in collaboration with Larry Young and Arjen Boender (titers 3.0 × 10¹⁰ and 1.5 × 10¹⁰ genomic copies/μl for gRNA and Cas9 AAVs, respectively). Retrograde rAAVs (retro-rAAV-hSyn-GFP, retro-rAAV-Ef1a-Flp, titer ≥ 7 × 10¹² vg/mL) were purchased from Addgene. The plasmid for GRAB-OT1.9 was obtained from the Yulong Li laboratory.

### Stereotactic Injections

Rats were anesthetized with either isoflurane (4% in 1 L/min O₂) or ketamine/xylazine (75 mg/kg and 5 mg/kg, respectively). Viral solutions (300 nl per hemisphere) were delivered at 150 nl/min using glass micropipettes. Injection coordinates (mm from bregma) were: PVN (AP −1.8, ML ±0.3, DV −8.0), SON (AP −1.8, ML ±1.2, DV −9.25), ILC (AP +3.0, ML ±0.5, DV −4.6), BLA (AP −3.0, ML ±5.0, DV −8.4 to −9.0), and NAc (AP +1.2, ML ±1.0, DV −7.8 to −8.2). Following all experiments, injection sites and transgene expression were confirmed histologically.

### Behavioral Tests

Behavioral tests were performed in an odor-resistant square arena (60 × 60 × 60 cm) under dim light (<20 lux, SO 200K lux-meter, Sauter). Rats were habituated to the arena for 15 min the day before testing, and the arena was cleaned with 80% ethanol between trials. Experimental and stimulus rats were housed separately.

#### Free Social Interaction

Experimental rats were paired with novel, age-and weight-matched conspecifics for 5 min. Animals were placed in diagonally opposite corners at trial start. For optogenetic experiments, 470 nm light (10 mW/mm², 30 Hz, 10 ms pulses) was delivered during the first 2 min of the session. For chemogenetic activation, the DREADD agonist CNO (3 mg/kg) was injected 40 min prior to testing. Behaviors (anogenital/non-anogenital sniffing, following, approach, toy investigation) were manually scored using BORIS software (https://www.boris.unito.it) by two independent, blinded experimenters. Discrepancies were resolved by consensus.

#### Food vs. SocialPpreference Test

The arena was equipped with triangular corner dividers containing mesh fronts, allowing indirect investigation of either an unfamiliar juvenile female rat (3–4 weeks) or standard food pellets. Each 5-min test session followed a 15-min habituation period. The test was repeated after 24 h of food deprivation. Investigation time was analyzed using Ethovision XT v11.5 (Noldus) by measuring nose position within predefined zones. The following parameters were quantified: total investigation time, number of bouts, and mean bout duration (a bout was defined as each re-entry after leaving the zone). Control experiments used the same setup, but only food was presented in one corner, with the other left empty.

### *Ex Vivo* Electrophysiology and Imaging

#### Slice Preparation

Brain slices were prepared 3–6 weeks after viral injections. For experiments 1–4 (1: TGOT application in current clamp, 2: TGOT application in voltage clamp, 3: paired-patch recordings, 4: optogenetic stimulation of OTR⁺ neurons with recordings from OTR⁻ pyramidal cells), slice preparation and visualization followed established methods (Stuart, Dodt and Sakmann 1993, Markram, Lübke et al. 1997). For experiment 5 (GRAB-OT1.9 sensor imaging combined with optogenetic stimulation of OT axons in the ILC), slice preparation followed Iwasaki, Lefevre et al. (2023). For experiment 6 (optogenetic stimulation of OTR⁺ interneurons with recordings from BLA-projecting pyramidal neurons), rats were anesthetized with isoflurane, decapitated, and brains rapidly removed into chilled, carbogenated ACSF containing (in mM): 92 choline chloride, 2.1 KCl, 1.2 NaH₂PO₄, 30 NaHCO_3_, 20 HEPES, 25 glucose, 5 Na-ascorbate, 2 thiourea, 3 Na-pyruvate, 5 N-acetyl-L-cysteine, 10 MgSO_4_, and 0.5 CaCl₂, pH 7.3–7.4, 290–310 mOsm. Coronal vmPFC sections (400 µm) were cut using a Leica VT1200 vibratome, incubated for 10 min at 32 °C in the same solution, and then transferred to recovery ACSF containing (in mM): 92 NaCl, 2.1 KCl, 1.2 NaH₂PO₄, 30 NaHCO₃, 20 HEPES, 10 glucose, 5 Na-ascorbate, 2 thiourea, 3 Na-pyruvate, 5 N-acetyl-L-cysteine, 1.3 MgSO₄, and 2.4 CaCl₂, pH 7.3–7.4, 290–310 mOsm, where they were kept for at least one hour before recording. After recordings, selected slices were fixed overnight in 4% PFA at 4 °C and immunostained to examine recorded neurons. Sections were incubated with chicken anti-GFP (1:10,000), rabbit anti-RFP (1:2,000), and Streptavidin-iFluor750 (1:4,000) in 0.5% Triton X-100/PBS for 72 h at 4 °C, washed in PBS, and then incubated with Alexa Fluor–conjugated secondary antibodies (1:2,000) for 24 h. Sections were mounted with Mowiol and imaged on an Olympus VS200 slide scanner and Leica Stellaris 5 confocal microscope.

#### Patch-Clamp Recordings

Slices were transferred to a recording chamber on a Nikon Eclipse FN1 microscope and perfused at 2 ml/min with carbogenated ACSF (32 °C). For experiments 1–4 the ACSF contained (in mM): 125 NaCl, 25 NaHCO₃, 2.5 KCl, 1.25 NaH₂PO₄, 2 CaCl₂, 1 MgCl₂, and 25 glucose, pH 7.2, 290–300 mOsm. For experiment 5 the ACSF contained (in mM): 118 NaCl, 25 NaHCO₃, 3 KCl, 1.2 NaH₂PO₄, 2 CaCl₂, 1.3 MgSO₄, and 10 glucose, pH 7.4, 290–300 mOsm.

Patch pipettes (5–7 MΩ) were filled with experiment-specific internal solutions. For experiments 1 and 4 the solution contained (in mM): 144 K-gluconate, 4 KCl, 4 Mg-ATP, 10 phosphocreatine, 0.3 GTP, and 10 HEPES, pH 7.3 with KOH, 293 mOsm. For experiment 2 it contained 144 Cs-gluconate, 4 CsCl, 4 Mg-ATP, 10 phosphocreatine, 0.3 GTP, and 10 HEPES, pH 7.3 with CsOH, 293 mOsm. For experiment 3 it contained 110 K-gluconate, 30 KCl, 8 NaCl, 4 Mg-ATP, 10 phosphocreatine, 0.3 GTP, and 10 HEPES, pH 7.3 with KOH, 293 mOsm. For experiment 6 it contained 145 K-gluconate, 2 MgCl₂, 4 Na₂ATP, 0.4 Na₃GTP, 5 EGTA, and 10 HEPES, pH 7.3 with KOH, 290–300 mOsm, and 0.05% biocytin for post-hoc neuron identification.

Whole-cell configuration was achieved by gentle negative pressure. Signals were recorded with an EPC 10 USB amplifier (HEKA), low-pass filtered at 2.9 kHz, digitized at 50 kHz.

#### Experimental Protocols and Analysis

For experiment 3 presynaptic neurons were stimulated with a 10 Hz train of three suprathreshold current pulses every 7 s, and postsynaptic responses were averaged over 50–100 sweeps. The success rate of synaptic coupling was calculated as the inverse (in %) of the number of neurons tested before a connection was found. In experiment 2 spontaneous EPSCs and IPSCs were recorded at −90 mV (E_GABA_) and 0 mV (E_AMPA_), respectively, and detected using a Wiener deconvolution algorithm in MATLAB (deconvwnr function) with τ_onset/τ_offset set to 4/15 ms for IPSCs and 2.5/10 ms for EPSCs. Peaks were identified in a 50 ms sliding window using dynamic and absolute thresholds, and PSC features were extracted from traces around each peak. In experiment 6 only layer 2/3 ILC pyramidal neurons projecting to the BLA were analyzed. Cells were voltage-clamped at −60 mV and stimulated with ten 470 nm light pulses (10 ms, ∼10 mW) at 30 s intervals, which were later averaged.

Data were analyzed with IGOR PRO (Wavemetrics), Signal (CED), and MATLAB. Electrophysiological results are reported as mean ± SD. Cumulative distributions were compared with the Kolmogorov– Smirnov test.

#### GRAB OT1.9 Oxytocin Sensor Imaging

Coronal brain slices containing the ILC were allowed to recover for at least one hour in oxygenated ACSF at room temperature after preparation. Expression of rAAV-OTp-ChrimsonR-mScarlet in the PVN and SON was confirmed using an X-Cite 110LED illumination system (XT-640-W). GRAB-OT1.9 imaging was performed on a Zeiss Axio Examiner microscope equipped with a CREST X-Light spinning disk confocal unit and a 20× water-immersion objective (NA 1.0). Images were acquired at 2 Hz using an optiMOS sCMOS camera (QImaging) controlled by MetaFluor software (v7.8.8.0). The OT1.9 sensor was excited at 475 nm (30 ms illumination), and ILC-projecting OT fibers were stimulated with 635 nm light at 30 Hz, 10 ms pulse width for 30 s after a 5 min baseline. Slices were then incubated in oxygenated ACSF with 1 µM Atosiban for ≥1 h before the second recording.

Fluorescence intensity was quantified using Fiji (ImageJ) and processed with a custom Python script. Signals were corrected for photobleaching by fitting a mono-exponential or polynomial decay model, and light-evoked OT release was quantified as the area under the curve (AUC) in a 50–350 s window post-stimulation, as previously described (Iwasaki, Lefevre et al. 2023).

### *In Vivo* Fiber-Optic Experiments

#### Optic Fiber Implantation

For *in vivo* optogenetic experiments, rats were bilaterally injected with rAAV-OTp-Venus or rAAV-OTp-ChR2-mCherry into the hypothalamic nuclei (SON, PVN) or with rAAV-Ef1a-DIO-ChR2-mCherry into the ILC, followed by bilateral optic fiber implantation above the ILC. Fibers (Ø 200 μm, NA 0.39, Thorlabs CFMC12L10) were cut to 5.5 mm length.

For *in vivo* fiber photometry, rats received bilateral injections of rAAV-Ef1a-DIO-GCaMP6s into the ILC and bilateral optic fiber implants (Ø 400 μm, NA 0.50, 5.5 mm, Doric Lenses) above the ILC.

Implantation coordinates for the ILC were AP +3.1 mm, ML ±1.52 mm, DV −4.75 mm, with a 10° angle to allow connection of patch cables.

#### Optogenetic Stimulation of OT Fibers

Two 473 nm blue lasers (DreamLasers, Shanghai) were coupled to patch cables (Thorlabs RJPFF2-FC/PC) and controlled by a Master-8 pulse generator. Light was delivered at 10 mW/mm², 30 Hz, 10 ms pulses for 120 s, as described previously (Knobloch, Charlet et al. 2012).

#### Fiber Photometry of ILC OTR Neurons

Rats were recorded for 10 min in a square open field arena. After 5 min habituation, an unfamiliar female rat or toy was introduced. Each rat underwent multiple sessions (44 total recordings across four animals). Data were acquired using a Doric photometry system and Doric Neuroscience Studio v6.1 software with 470 nm (Ca²⁺ signal) and 405 nm (isosbestic) excitation at 20 Hz and 0.4 mW output power. Videos were recorded in parallel for behavioral annotation.

Raw signals were exported and preprocessed using the pMat toolbox (Bruno, O’Brien et al. 2021). High-frequency noise was removed by smoothing, and ΔF/F was calculated as (calcium signal – scaled isosbestic signal) / scaled isosbestic signal. The ΔF/F traces were then Z-scored relative to the mean and standard deviation of the baseline. For each behavioral event (anogenital sniffing, non-anogenital sniffing, toy sniffing), we compared the normalized signal averaged over 1 s before event onset to the signal averaged over the first 1 s after onset.

Statistical analysis was performed in R (Version 4.0.4; R Core Team, 2021) using RStudio (Version 1.4.1106, PBC; Team R, 2019). Linear mixed-effects models were fitted using the *lme4* (Douglas Bates, Bolker and Walker 2015) and *lmerTest* (Kuznetsova, Brockhoff and Christensen 2017) packages, with type III ANOVA performed using Satterthwaite’s method. Post hoc Tukey’s multiple comparisons were conducted with the emmeans package (Lenith 2023), and the Kenward–Roger approximation for denominator degrees of freedom. Model assumptions were verified by inspection of residual plots.

The models compared Z-scored ΔF/F values before vs. during each behavioral event in a paired manner, with trial included as a random effect nested within session and sessions nested within animal. When recordings were performed bilaterally, hemisphere was added as a cross-nested random effect. Fixed factors included time in relation to trial onset (before vs. during the behavior), and in the case of comparison of active and received behaviors also behavior type. The model for the behavior towards the toy was fitted separately, as these were recorded from separate sessions.

### Histology

Rats were deeply anesthetized and transcardially perfused with PBS followed by 4% paraformaldehyde (PFA). Brains were post-fixed overnight in 4% PFA at 4 °C, sectioned coronally at 50 µm on a vibratome (Leica VT1000S), and collected into series spanning the rostrocaudal extent of the hypothalamic OT system and the ILC. Free-floating sections were processed using established protocols. Primary antibodies included anti-OT (PS38, mouse, 1:2000, gift of H. Gainer), anti-DsRed (Clontech 632397, rabbit, 1:1000), anti-GFP (Abcam ab13970, chicken, 1:1000), anti-cFos (Cell Signaling 9F6, rabbit, 1:500), anti-NeuN (Merck MAB377, mouse, 1:1000), anti-Ankyrin (Millipore MABN466, mouse, 1:1000), anti-Parvalbumin (Abcam ab427, rabbit, 1:5000), anti-Somatostatin (BMA T-4103, rabbit, 1:3000), anti-Calbindin (SWANT CB-300, mouse, 1:3000), and anti-Calretinin (SWANT 6B3, mouse, 1:5000). Sections were incubated with primary antibodies in PBS with 0.5% Triton X-100 for 48–72 h at 4 °C, rinsed, and incubated with secondary antibodies (1:1000; Alexa 488, Alexa 594, Alexa 647, or Cy3-conjugated donkey secondaries; Jackson ImmunoResearch or Invitrogen) for 24 h. Sections were mounted in Mowiol for imaging. Whole-section scans were acquired on an Olympus SLIDEVIEW VS200 slide scanner for overview images and verification of viral injection sites. Images of individual neurons, co-labeled markers, and OT fiber proximity were captured on a Leica Stellaris 5 confocal microscope in Z-stack mode.

#### Fiber Quantification Analysis

Densities of OT-positive fibers were quantified in serial sections collected between Bregma +3.72 and +2.52, corresponding to the rostral and caudal borders of the ILC (n = 11 sections from 5 animals). Anatomical borders between mPFC subdivisions were defined based on the Paxinos Rat Brain Atlas and previous cytoarchitectonic studies (Kolk and Rakic 2022). Whole-brain section scans containing Cg1, PrL, and ILC were captured with an Olympus VS200 slide scanner at 20× magnification and stitched into VSI image files. Fiber densities were quantified by counting crossing points of OT fibers in digitized images overlaid with a non-destructive grid of lines (Mamounas, Altar et al. 2000) in ImageJ. Crossing points were averaged per animal, normalized to region area, and presented as fibers/mm².

#### Neuron Quantification Analysis

For each animal, one of four series of 50 µm sections through the ILC was analyzed to quantify OTR-expressing neurons. Two analyses were performed: (1) the proportion of inhibitory vs. excitatory OTR-expressing neurons across the ILC, and (2) co-expression of cytochemical markers. Neurons were manually counted in ImageJ using touch-count mode, averaged, and presented as percent of total OTR-expressing neurons.

#### Automated Image Analysis

Proximity of OT fibers to OTR neurons was quantified in Imaris (Bitplane, *Oxford Instruments*). Green (OTR⁺ interneurons) and red (OTR⁺ neurons) channels were surface-reconstructed using a grain size of 0.6 µm, sphere diameter 0.5 µm, with threshold and volume parameters adjusted per image (volumes >1.7–5.25 µm³ included for green, >10 µm³ for red). Blue-channel OT fibers were reconstructed using a grain size of 0.44 µm and included if >3.85–10 µm³. Fiber segments were manually identified and assigned ID numbers for proximity mapping. GFP-, cFos-, and NeuN-positive cells were quantified in Imaris (Spots function, 10 µm diameter, quality threshold 1.21–1.35). The ILC was segmented manually into layers 1, 2/3, and 5/6 based on anatomical landmarks, and neurons were assigned accordingly. OT fiber distribution across ILC layers was calculated from surface reconstructions with a surface detail of 0.89 µm, absolute intensity threshold 2.2–6, and minimum volume 70 µm³.

### Western Blot

Protein was extracted from OTR-Cre and WT rat brains using RIPA Lysis buffer, quantified with a BCA kit, separated on 4–12% NuPage Bis-Tris gels (Invitrogen), and transferred to nitrocellulose membranes. Membranes were blocked in 5% milk/TBS-T, incubated overnight at 4 °C with rabbit anti-oxytocin receptor (Thermo Fisher, 1:1000) and mouse anti-GAPDH (Abcam, 1:5000), followed by HRP-conjugated secondary antibodies. Bands were visualized by chemiluminescence and quantified with ImageJ.

### Multiplex Fluorescent *In Situ* Hybridization

Expression profiles of OTR neurons in the ILC were characterized using RNAscope™ HiPlex (Advanced Cell Diagnostics, Hayward, CA, USA) on fresh-frozen 16 µm sections (two ILC slices per brain, five animals) as previously described (Szlaga, Sambak et al. 2022). The following probes were used: vGlut1 (Rn-Slc17a7-T1, cat. no. 317001-T1), Cre (CRE-T2, cat. no. 312281-T2), vGAT1 (Rn-Slc32a1-T3, cat. no. 424541-T3), vGlut2 (Rn-Slc17a6-T5, cat. no. 31701-T5), and OTR (Rn-Oxtr-T6, cat. no. 483671-T6). Imaging was performed on a Zeiss Axio Imager M2 equipped with an automated z-stage and Axiocam 503 mono camera. Images were processed with Zen (3.1 blue edition and 3.0 SR black edition, Zeiss), CorelDraw 2020 (Corel), ImageJ (Schneider, Rasband and Eliceiri 2012), and HiPlex Image Registration Software v1.0 (ACD). OTR mRNA-positive cells were identified by DAPI-stained nuclei or cell-like mRNA distribution and manually counted in ImageJ (Cell Counter), following Iwasaki, Lefevre et al. (2023). Total neuronal populations were quantified in QuPath (Bankhead, Loughrey et al. 2017) with the following parameters: detection[Channel 2]-vGlut1 75, detection[Channel 6]-vGAT 20, doSmoothing TRUE, splitByIntensity TRUE, splitByShape TRUE, spotSizeMicrons 2, minSpotSizeMicrons 0.15, maxSpotSizeMicrons 6, includeClusters TRUE.

### Statistics

An Excel file containing all statistical tests and results is provided as **Supplemental Document 1**. For each dataset, normality was assessed before selecting the appropriate test. Parametric tests (paired or unpaired two-tailed t-tests, two-or three-way ANOVA) were used when assumptions were met; otherwise, non-parametric tests (Wilcoxon signed-rank, Mann–Whitney U) were applied. For repeated-measures or unbalanced designs, linear mixed-effects models were used. Unless stated otherwise, α was set at 0.05 and post hoc comparisons were corrected for multiple testing. In figures, asterisks (*) indicate significant main effects and hash symbols (#) indicate significant post hoc effects. Fiber-photometry data were analyzed using linear mixed-effects models as described in the section Fiber Photometry of ILC OTR Neurons.

## DATA AVAILABILITY

Source Data are provided with this paper. We provide two Supplemental Documents with this publication:

**Supplemental Document 1** includes statistical test results and details for all performed tests, divided on individual Excel sheets per Figure.

**Supplemental Document 2** includes all plotted data in individual Excel sheets.

## CODE AVAILABILITY

The python script used for analysis of OT1.9-GRAB-sensor fluorescent signal is made available with this publication. Additionally, the R code for the statistical analysis of fiber photometry data is made available with this publication. Any readily available software and packages we utilized are detailed in the method section.

## EXTENDED DATA

**Extended Data Figure S1.**
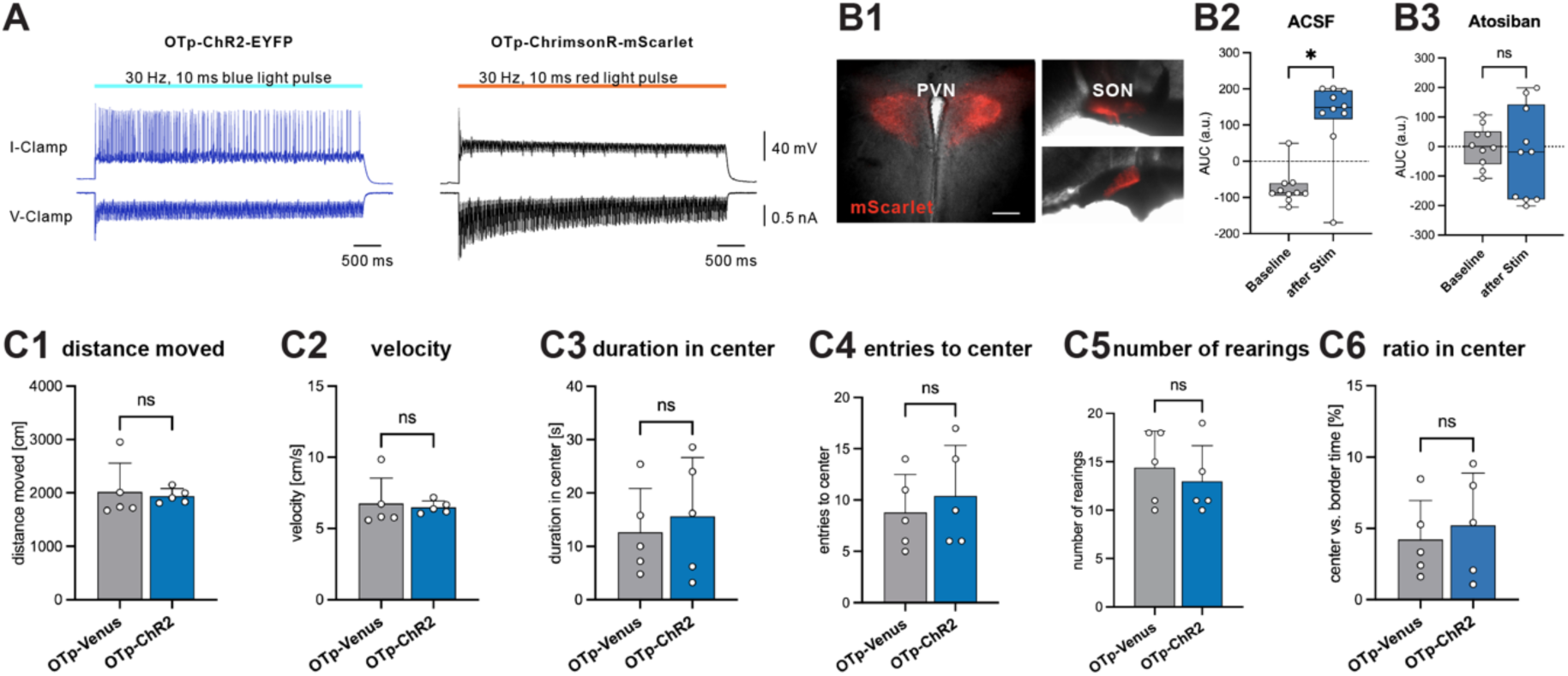
**A** Functional verification of OTp-ChR2 and OTp-ChrimsonR through patch-clamp recordings. **B1** Representative OTp-ChrimsonR-mScarlet expression in the PVN and SON. **B2** AUC for OT-sensor signal before and after stimulation of OT fibers in the ILC with red light. **B3** AUC for OT-sensor signal before and after stimulation of OT fibers in the ILC with red light after bath application of an OTR-antagonist (Atosiban) (n=10 sections). **C** Exploratory behavior in the open field after optogenetic activation of OT axons in the ILC. **C1** Distance moved and **C2** velocity in the open field, a control for locomotor effects, remain unchanged by optogenetic manipulation of OT axons in the ILC. **C3** Duration in center and **C4** entries to center, a control for anxiety-like behavior, remain unchanged by optogenetic manipulation. **C5** Number of rearings as a non-social behavior is not affected by optogenetic manipulation (n=5 animals per group). **C6** The percentage of time spent in the center of the arena remains unchanged. Statistical significance is indicated as * p<0.05, ** p<0.01, *** p<0.001, **** p<0.0001. Statistical significance of post-hoc tests is indicated as # p<0.05, ## p<0.01, ### p<0.001, #### p<0.0001. Error bars show standard deviation. For details on statistical tests please refer to Supplemental Document 1.

**Extended Data Figure S2.**
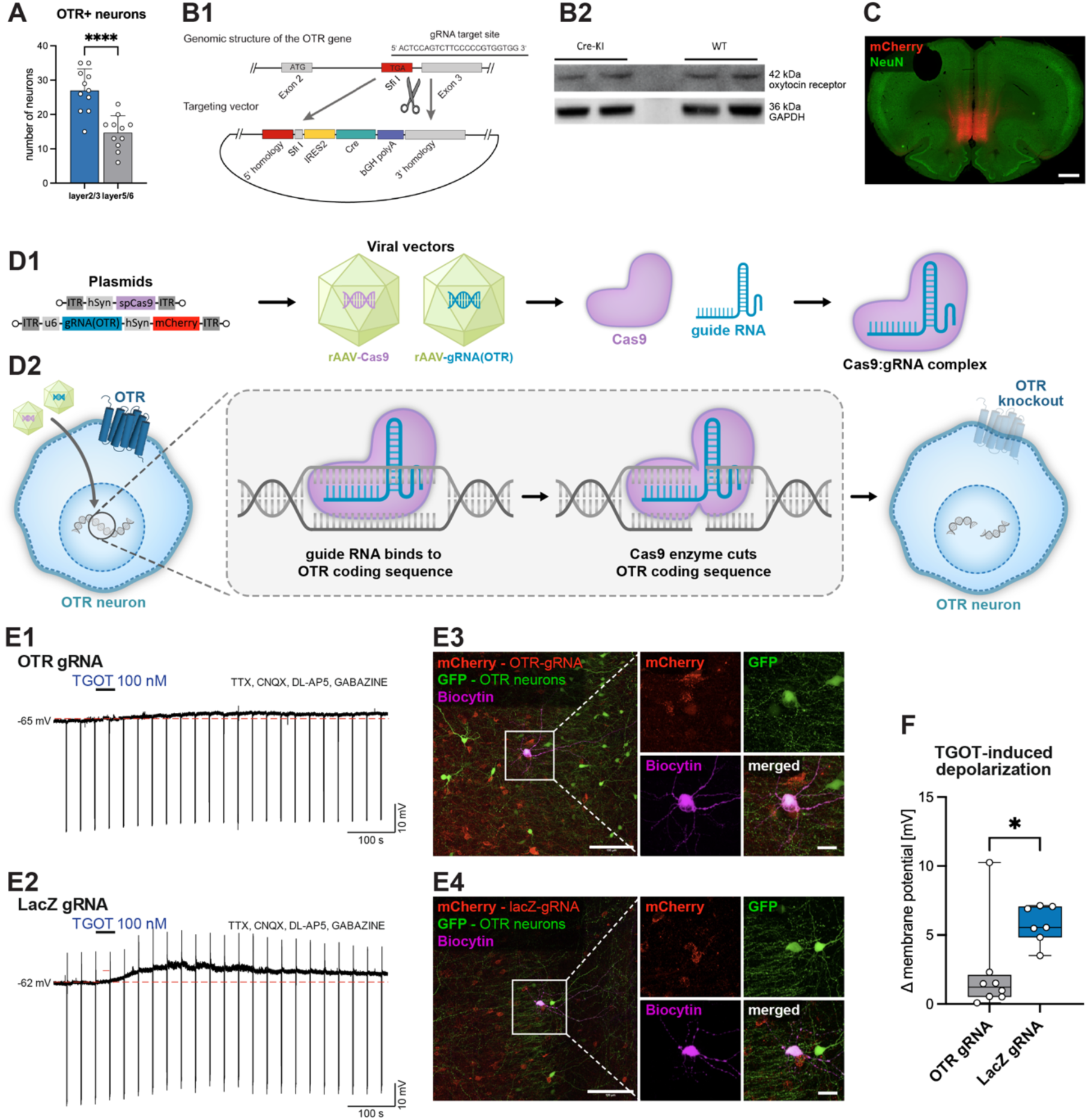
**A** More OTR^+^ neurons are found in layer 2/3 of the ILC, compared to layer 5/6. **B1** Generation of the transgenic OTR-Cre rat line utilizing CRISPR/Cas9. B2 Western blot analysis of the OTR-levels in both transgenic rat line (Cre-KI) and wildtype (WT). **C** Injection site of the Cas9/gRNA virus system in the ILC. **D** Mechanism of OTR depletion in neurons utilizing a viral CRISPR/Cas9 approach. **D1** Two viral vector plasmids are used to produce rAAVs expressing either the Cas9 enzyme or a specific guide RNA for OTR. **D2** After injection of the mixture of both viruses into the ILC, the Cas9:gRNA complex splices the OTR coding sequence in all neurons and leads to a partial functional OTR knockout, while the neurons themselves remain viable. As a control, a guide RNA for lacZ is used, as no unspecific splicing is expected. **E** Representative traces of a whole-cell patch clamp recordings (current clamp, zero current mode) obtained from an OTR^+^ ILC neuron transfected with AAVs carrying **E1** Cas9 and OTR gRNA or **E2** Cas9 and LacZ gRNA. A TGOT-induced depolarization in the control condition (LacZ) and its absence in the OTR loss-of-function was observed. Downward deflections represent voltage responses of the recorded neuron to hyperpolarizing current injections. **E3** Series of fluorescent projection images of biocytin-filled OTR^+^ ILC neurons transfected with AAVs carrying Cas9 and OTR gRNA or **E4** Cas9 and LacZ gRNA. **F** TGOT-induced depolarization is significantly reduced in the OTR^+^ ILC neurons transfected with AAVs carrying Cas9 and OTR gRNA (n=8 neurons) compared to the control condition (n=7 neurons). Statistical significance is indicated as * p<0.05, ** p<0.01, *** p<0.001, **** p<0.0001. Statistical significance of post-hoc tests is indicated as # p<0.05, ## p<0.01, ### p<0.001, #### p<0.0001. Error bars show standard deviation. For details on statistical tests please refer to Supplemental Document 1.

**Extended Data Figure S3.**
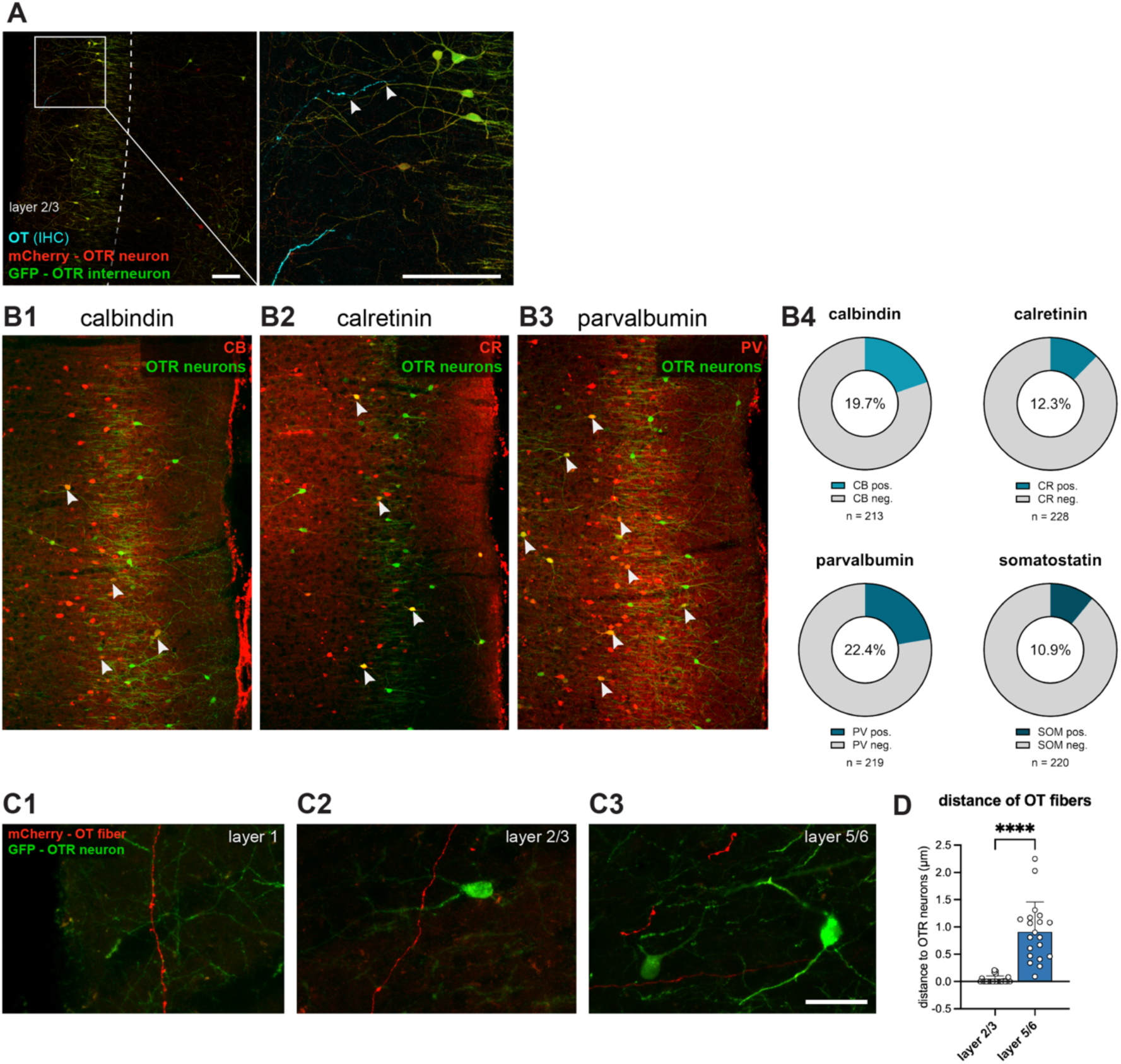
**A** Alternative representative scan to Figure 3 **B2**. Scalebar 100µm. **B** Heterogenous OTR interneuron population in the ILC. **B1-C3** Representative images of ILC sections previously injected with a Cre-dependent rAAV expressing GFP in OTR neurons and counterstained for various interneuron markers: calbindin (CB), calretinin (CR) and parvalbumin (PV). Scalebar. **B4** Quantitative analysis of the OTR interneuron population; both calbindin or parvalbumin are expressed in about 20% of OTR interneurons in the ILC, while calretinin or somatostatin are found in 10% of interneurons (n=4 animals). Scalebar 150µm. C OT fibers in close proximity to OTR^+^ neurons. OT fibers labelled by viral injection of rAAV-OTp-mCherry in the PVN and SON are found **C1** close to the midline in layer 1, **C2** in close proximity to an OTR^+^ interneuron in layer 2/3, and C3 in close proximity to a pyramidal neuron in layer 5/6. Scalebar 40µm. D Automated, high-throughput analysis of the distance of OT fibers to OTR neurons in the ILC. Fibers are closer to neurons in layer 2/3 compared to layer 5/6 (n=24 neurons and n=20 neurons, sections obtained from 4 animals). Statistical significance is indicated as * p<0.05, ** p<0.01, *** p<0.001, **** p<0.0001. Statistical significance of post-hoc tests is indicated as # p<0.05, ## p<0.01, ### p<0.001, #### p<0.0001. Error bars show standard deviation. For details on statistical tests please refer to Supplemental Document 1.

**Extended Data Figure S4.**
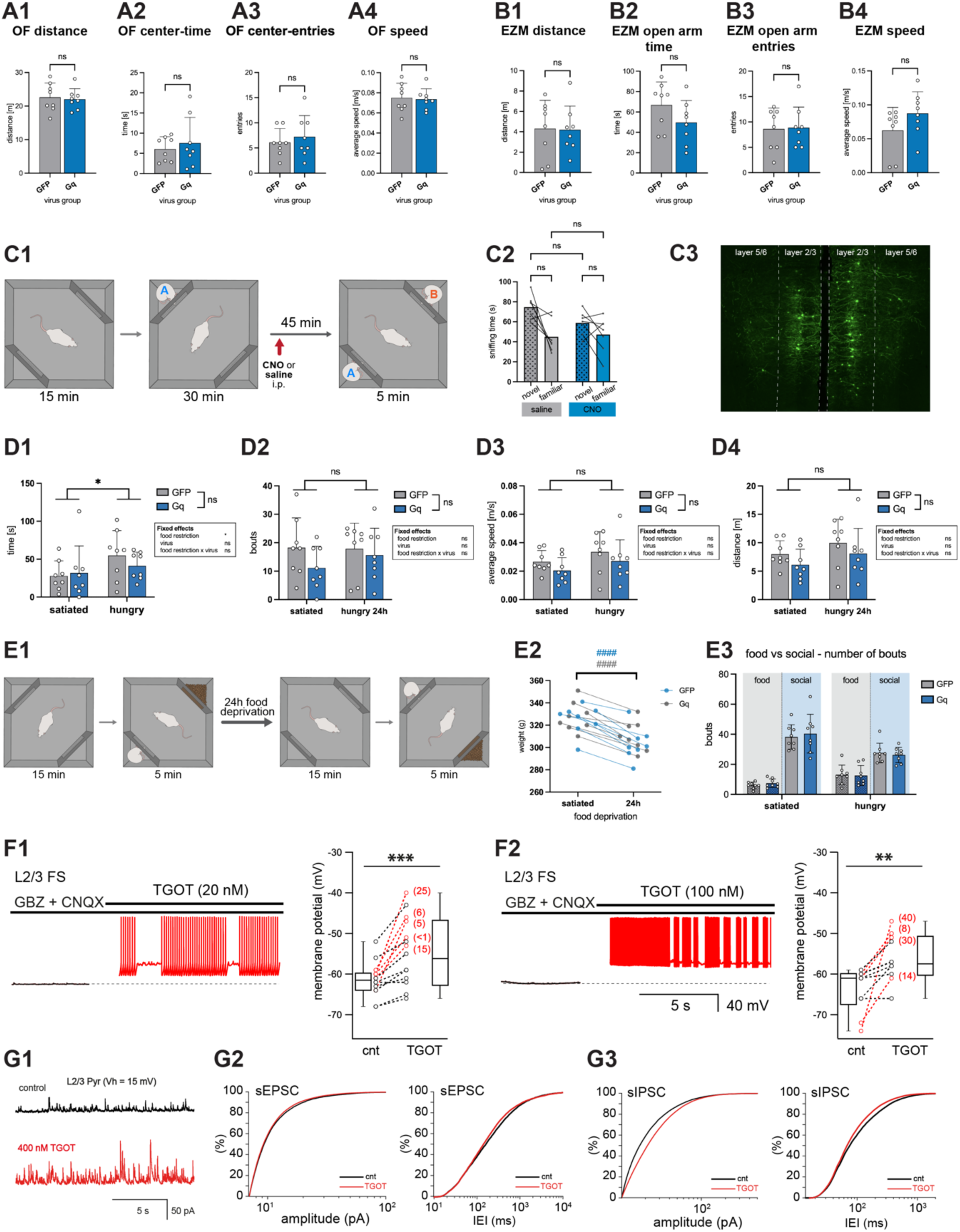
**A** Exploratory behavior in the open field (OF) after chemogenetic activation of OTR^+^ interneurons in the ILC. **A1** Distance travelled in the OF, **A2** Time in the center of the OF, **A3** number of entries into the OF, and **A4** average speed in the OF (n=8 animals). **B** Exploratory behavior in the elevated zero maze (EZM). **B1** Distance travelled in the EZM, **B2** Time spent in the open parts of the EZM, **B3** number of entries into the open parts of the EZM, and **B4** average speed in the EZM (n=8 animals). **C** Social novelty preference paradigm. **C1** Paradigm timeline: animals were habituated to an open field arena with two empty corner compartments for 15 minutes, before placing an unknown female conspecific (animal A) into one corner for a 30 min familiarization session; after a 45 min break, both the familiar (A) and a novel (B) conspecific rat were placed in opposite corners, in order to test social novelty preference. CNO was administered 40 min before the start of the 5 min test session. Corners were placed randomly and altered between sessions and animals. **C2** No effect of CNO administration was observed compared to saline administration (n=7 animals). **C3** Representative injection site of rAAV-DLX-DIO-Gq-GFP for the food vs. social paradigm and social novelty preference paradigm, as verified for all experimental animals post-hoc. Scalebar 500µm. **D** Food interest. Animals were placed in an OF with food pellets behind a mesh in one corner. **D1** The time spent investigating the food was only influenced by hunger, but not chemogenetic activation of OTR^+^ interneurons. **D2** Number of bouts while investigating food in the OF. **D3** Average speed in the OF as well as **D4** distance travelled in the OF (n=8 animals per group). **E** Food vs. social behavior paradigm. **E1** Paradigm timeline: animals were habituated to an open field arena with two empty corner compartments for 15 minutes, before placing an unknown female conspecific into one corner, and food into the opposite corner; after 24h of food deprivation the test was repeated. CNO was always injected 40 min before the start of the 5 min choice session. Corners were placed randomly and altered between sessions and animals. **E2** Weight loss of rats after 24h of food deprivation was verified for both groups. **E3** The number of bouts for food and social corners were unaltered (n=8 animals per group). F Representative traces of whole-cell patch clamp recordings (current clamp, zero current mode) and quantitative analysis of membrane potential changes in layer 2/3 fast-spiking interneurons upon bath application of either F1 20nM TGOT (n=14 cells, sections obtained from 4 rats; p=0.0005) or F2 100nM TGOT (n=10 cells, sections obtained from 4 rats; p=0.0097). **G** Analysis of EPSCs (n=37) and IPSCs (n=38) of layer 2/3 OTR negative pyramids after TGOT application (400nM) compared to baseline (cnt). The brain sections were obtained from 10 rats. **G1** Representative trace of IPSC recording. **G2** No effect on EPSC amplitude or kinetics was observed, however **G3** IPSC amplitude (and charge transferred) are increased (20.7 ± 2.3 vs. 23.6 ± 1.8 pA, p = 0.0017; 2173 ± 222 vs. 2657 ± 837 pC, p=0.0013, respectively). IEI: inter-event-interval. Statistical significance is indicated as * p<0.05, ** p<0.01, *** p<0.001, **** p<0.0001. Statistical significance of post-hoc tests is indicated as # p<0.05, ## p<0.01, ### p<0.001, #### p<0.0001. Error bars show standard deviation. For details on statistical tests please refer to Supplemental Document 1.

**Extended Data Figure S5.**
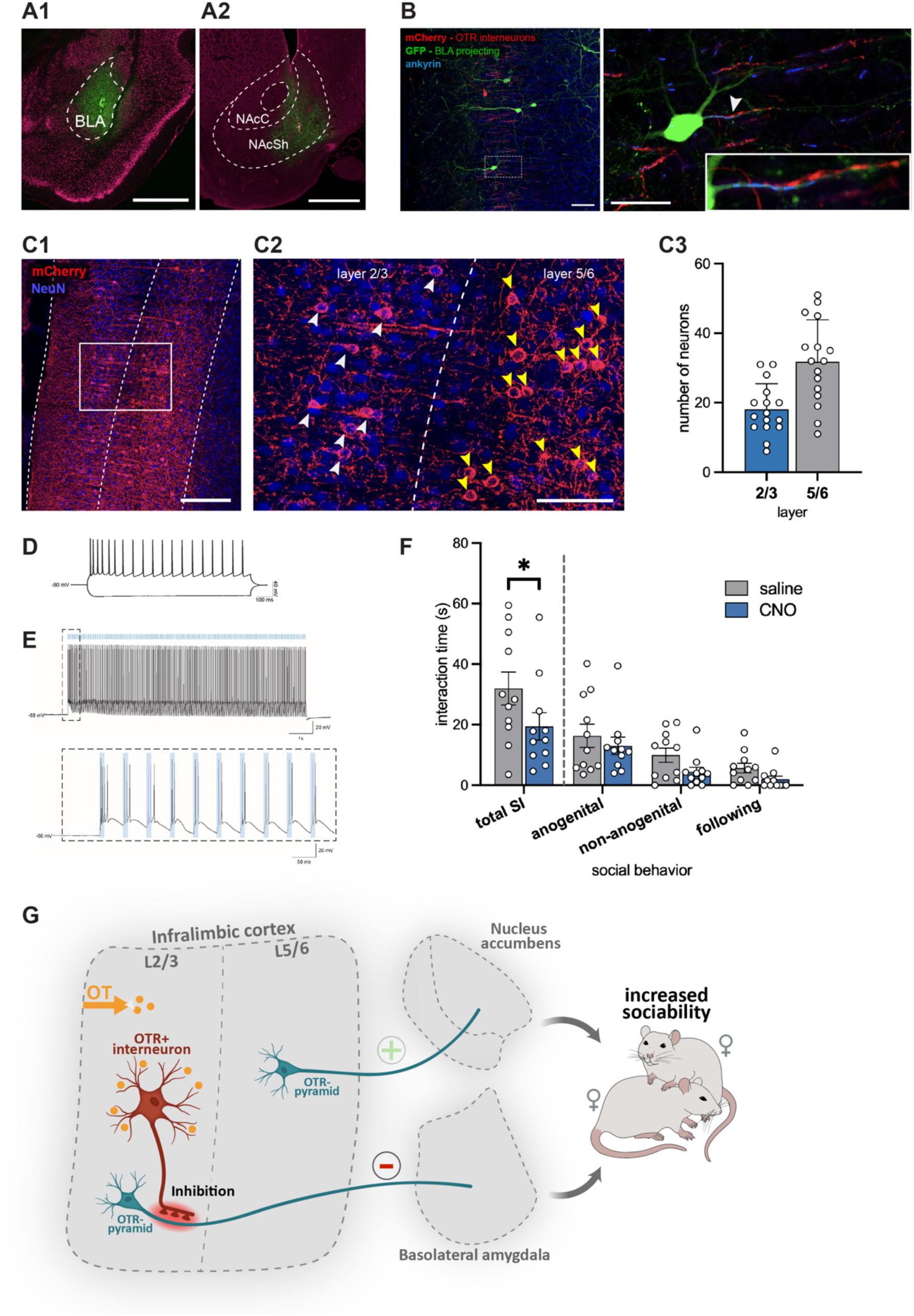
**A** Representative injection sites of retro-AAV-hSyn-GFP in the **A1** BLA or **A2** NAc. The position of the pipette tip during the injection is marked with retrobeads (white arrow). **B** Representative axo-axonic contact (white arrow) of OTR+ interneuron (red) on BLA-projecting pyramidal neuron (green). Scalebar 50μm overview, 25µm zoom. **C** Quantification of BLA-projecting neurons in layer 2/3 and layer 5/6 of the ILC. **C1** Overview of the ILC with BLA-projecting neurons in red (rAAV-FRT-Gq-mCherry in ILC, retro-rAAV-Flp in BLA) and NeuN staining (blue). Scalebar 250µm. **C2** BLA-projecting neurons in layer 2/3 (white arrows) and layer 5/6 (yellow arrows). Scalebar 100µm. **C3** Number of BLA-projecting neurons in layer 2/3 and layer 5/6 of the ILC. **D** Representative traces of a whole-cell patch clamp recordings (current clamp, zero current mode) obtained from a BLA-projecting ILC neuron showing spike frequency adaptation typical for pyramidal cells. **E** Representative traces of a whole-cell patch clamp recordings (current clamp, zero current mode) obtained from an OTR^+^ ILC interneuron expressing ChR2 and following to BL stimulation (20Hz, 10ms long pulses, ∼10mW). The right trace shows a zoom of the first 10 stimulations. **F** Working hypothesis. OT projections from the hypothalamus innervate the infralimbic cortex, where they specifically activate OTR^+^ interneurons in layer 2/3. These OTR^+^ interneurons, which show functional and anatomical properties of Chandelier interneurons, preferentially target and inhibit OTR^-^ pyramidal neurons projecting to the basolateral amygdala (BLA), decreasing neuronal activity in the BLA. Specific and targeted activation of this OTR^+^ neuronal population increases sociability and social preference in female rats. Statistical significance is indicated as * p<0.05, ** p<0.01, *** p<0.001, **** p<0.0001. Statistical significance of post-hoc tests is indicated as # p<0.05, ## p<0.01, ### p<0.001, #### p<0.0001. Error bars show standard deviation. For details on statistical tests please refer to Supplemental Document 1.

## Notes

### Competing Interest Statement

The authors have declared no competing interest.

### Summary of Updates

- utilisation of newly developed GRAB OT sensor to demonstrate that ex vivo optogenetic stimulation of OT axons in the ILC increased the GRAB OT sensor signal; this effect could be blocked by administration of the OT receptor (OTR) antagonist Atosiban - extended the quantitative analysis across several anatomical experiments: (1) expanded in situ hybridisation with additional animals (now n=5 rats); (2) quantified OT axon innervation and OTR expression in the ILC (subdivided by cortical layers and along the rostro-caudal axis); (3) quantified OTR+ Chandelier neurons; (4) subdivided the cFos analysis in the ILC by cortical layers; (5) retro-gradely traced and quantified basolateral nucleus of amygdala (BLA)-projecting pyramidal neurons - performed two new chemogenetic experiments: (1) behavioural role of inhibiting downstream projections to the BLA and (2) addressed experimental design flaws in the original food vs. social choice paradigm

